# Genome of the glacier alga *Ancylonema* and its insights into the evolution of streptophyte life on ice and land

**DOI:** 10.1101/2023.10.20.563300

**Authors:** Alexander M. C. Bowles, Tom A. Williams, Philip C. J. Donoghue, Douglas A. Campbell, Christopher J. Williamson

## Abstract

Contemporary glaciers and ice sheets are home to communities of streptophyte glacier algae that must balance their requirements for photosynthesis and growth with tolerance of extremes in temperature, desiccation and UV radiation. These same environmental challenges have been hypothesized as the driving force behind the evolution of land plants from streptophyte algal ancestors in the Cryogenian (720–635 million years ago). Here, we sequence, assemble and analyze the metagenome-assembled genome (MAG) of the glacier alga *Ancylonema nordenskiöldii* to investigate its adaptations to life in ice, and whether this represents a vestige of Cryogenian anydrophyte exaptations. Phylogenetic analysis confirms the placement of glacier algae within the sister lineage to land plants, Zygnematophyceae. The MAG is characterized by an expansion of genes involved in high irradiance and UV light tolerance, whilst lineage-specific diversification led to the novel screening pigmentation of glacier algae. We found no support for the hypothesis of a common genomic basis for adaptations to ice and to land in streptophytes. Comparative genomic analysis revealed that reductive genome evolution in the ancestor of Zygnematophyceae correlates with their reductive morphological evolution. This first genome-scale data for glacier algae supports an *Ancylonema*-specific adaptation to the cryosphere, as well as shedding light on the genome evolution of land plants and Zygnematophyceae.

## Introduction

The evolution of terrestrial flora (Embryophyta)^1^ transformed Earth’s continents, atmosphere and climate^2–6^, and promoted the diversification of multiple lineages spanning the tree of life^7,8^. The first land plants evolved from a streptophyte algal ancestor^1,9^, the last common ancestor of Anydrophyta, with time-scale analyses inferring a likely Neoproterozoic divergence between Embryophyta and its closest Zygnematophycean algal relatives. The youngest estimates fall within the Ediacaran period^10^, with others dating this divergence as far back as the Mesoproterozoic^11^. To make the fundamental transition from an aquatic to terrestrial environment required adaptations to tolerate extremes in temperature, desiccation and UV radiation^1,12–16^. It is likely that ancestral streptophytes possessed a suite of exaptations (pre-adaptations) available to be co-opted to these ends. Studying closely related extant streptophytes could provide significant insight into the genomic capacity of ancestral lineages and is therefore critical to unravelling the processes involved in land plant terrestrialisation^1,17^.

Zygnematophyte glacier algae live on the surfaces of contemporary glaciers and ice sheets, and proliferate in widespread algal blooms during summer melt seasons when sunlight and liquid water are available to power photosynthesis^18–22^. Given this climatically important, yet under-studied lifestyle, investigation into their adaptations to life in ice through genome-scale sequencing is imperative. To inhabit cryospheric environments, glacier algae must balance their requirements for photosynthesis and growth with tolerance of extremes in temperature, desiccation and UV radiation^18,21^, raising interesting parallels with the challenges faced by plants during their adaptation to life on land. Furthermore, one current hypothesis, as yet untested, suggests that plants moved onto land in the aftermath of global glaciations in the Cryogenian, whereby ice provided an intermediate habitat between water and land^23,24^. Thus, considering the niche of modern-day glacier algae, and their phylogenetic proximity to embryophytes, these species represent an important model system to explore adaptations to extreme conditions and the process of plant terrestrialisation.

Here, we describe and analyze the metagenome-assembled genome (MAG) of the glacier alga *Ancylonema nordenskiöldii* to investigate the genomic basis of its adaptations to life on ice and the process of plant terrestrialisation more broadly. Specifically, we explore whether modern day adaptations of glacier algae represent exaptations deriving from an anydrophyte ancestor, which would support the hypothesis that Cryogenian glaciations were a major driver of land plant evolution, or whether these adaptations emerged more recently within the glacier algal lineage. The MAG also enables comparative analyses of genome evolution in Zygnematophycaeae, to test the recent hypothesis^25,26^ that gene loss has dominated their evolutionary history.

## Results and Discussion

### The metagenome of the glacier alga Ancylonema nordenskiöldii

We present the first genome-scale data for glacier algae. A single glacier algal MAG was assembled using a combination of PacBio High-Fidelity (HiFi) long reads and Illumina short reads; in total, these were assembled into 9,979 contigs (25KB contig N50). The MAG was estimated at 170 megabases (Mb), consistent with previous Zygnematophyceae genomes (*Spirogloea muscicola*: 174Mb, *Mesotaenium endlicherianum*: 163Mb^27^, *Penium margaritaceum*: 4700Mb^28^). We predicted 19,593 protein-coding genes in the glacier algal MAG whilst BUSCO^29^ analysis (Eukaryota_odb10) suggested a genome completeness of 86.7% (Supplementary Information 1). This set of protein-coding genes was used for downstream analyses.

Current knowledge based on morphology and amplicon sequencing data (18S and rbcL) recognizes two species of glacier algae, the filamentous *Ancylonema nordenskiöldii* and the unicellular *Ancylonema alaskanum*^24^. Samples for the present study were collected in 2020 from Morteratsch glacier, Switzerland; a site known to harbor both glacier algal species, though with a dominance (70% relative abundance) of *A. nordenskiöldii*^30^. Reciprocal BLAST analysis of glacier algal 18S and rbcL marker genes against our contigs identified our MAG as *A. nordenskiöldii* (Supplementary Data 1-4). Additional analysis of GC content and kmers did not identify a bimodal distribution of contigs (Supplementary Information 2 & 3). Duplicated BUSCO genes were mostly involved in enzymatic processes, suggesting proliferation of metabolism in a single organism, rather than the presence of multiple related organisms in the sample (Supplementary Data 5). Therefore, we concluded that the MAG contained material from a single species, *A. nordenskiöldii*, and proceeded conservatively with downstream analyses on that basis. While we utilised a metagenomic approach due to lack of viable cultures, chromosome scale assemblies will be required to fully investigate the lives of glacier algae. Attempts to culture glacier algae have very recently come to fruition^31,32^, opening up the opportunity for complete genome characterisation in the future.

Phylogenetic analysis placed our *A. nordenskiöldii* MAG within the Zygnematophyceae, the sister group to land plants, with which they comprise the group Anydrophyta (Figure 1). Further analysis placed Anydrophyta within the Streptophyta, with Chlorophyta sister to this latter group (Supplementary Information 4). These results were consistent with previous analyses of Zygnematophyceae^25^, Streptophyta^9^ and green plant evolution^4^. Within the Zygnematophyceae, *A. nordenskiöldii* is placed within the Zygnematales (Figure 1). As discussed, morphological analysis has classified glacier algae as the filamentous *A. nordenskiöldii* and the unicellular *A. alaskanum* (previously *Mesotaenium berggrenii*)^24,33^. Our analysis of genome-scale data indeed placed glacier algae separate from *Mesotaenium* (Figure 1), highlighting the need for re-classification of *Mesotaenium* species as well as species in the genus *Cylindrocystis*.

**Figure 1.**
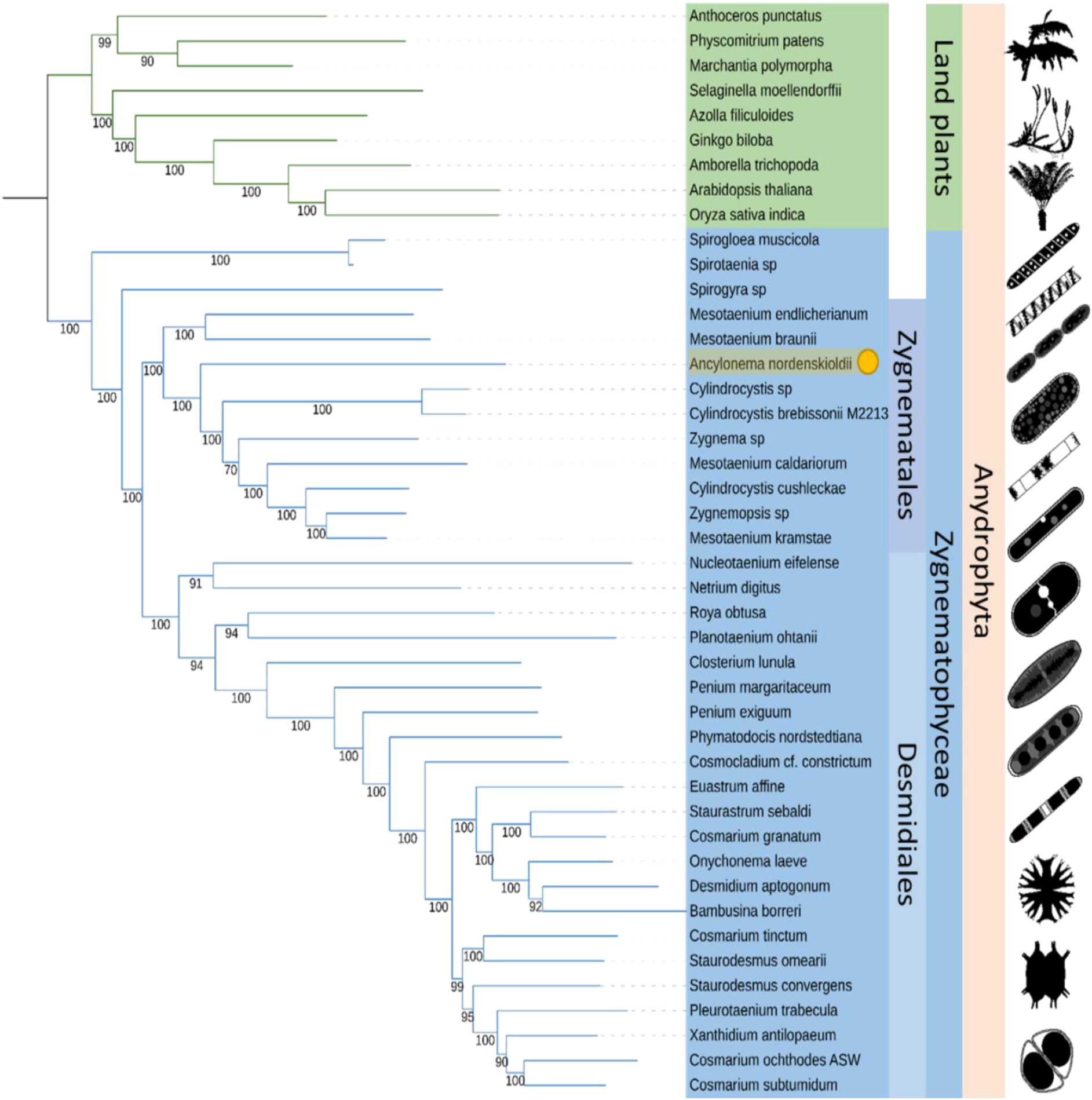
Phylogenetic analysis of *Ancylonema nordenskiöldii.* **A,** Phylogenetic analysis of the placement of *Ancylonema* within the Anydrophyta, based on a concatenated alignment of 543 genes. Taxa are colour coded: Land plants (green), Zygnematophyceae (blue). *Ancylonema nordenskiöldii* highlighted in yellow. Silhouettes are sourced from Phylopic (Supplementary Data 48).

### Glacier algal adaptations to life in ice

Comparative genomic analysis combining the *A. nordenskiöldii* MAG with previously published data indicated that glacier algae adapted to the cryosphere through lineage-specific diversification of existing genetic pathways (Figure 2a, b & c, Supplementary Data 6-15). The 19,593 protein coding genes from our *A. nordenskiöldii* MAG were clustered into 6,242 orthogroups (OGs) using OrthoFinder^34^. Gene ontology (GO) analysis of 2184 expanded gene families, identified with CAFE^35^, revealed functions associated with water transport (e.g. lipid localisation), protein repair (e.g. autophagy, PSII associated light-harvesting complex), response to abiotic stimulus (e.g. response to high light intensity, UV, radiation), wax metabolic process (e.g. lipid biosynthesis and modification) and plant-type cell wall modification (e.g. chloroplast organisation; Figure 3b, Supplementary Data 6, 8 & 12, Supplementary Information 5). The function of these expanded gene families suggested these algae gained adaptations to biotic and abiotic stressors, principally to high-light and UV stress, consistent with the known high-light environment inhabited by glacier algae^21^. This was likely achieved through duplication of genes involved in light sensing and photodamage repair, potentially from UV irradiation. Gene gain was low (276 genes gained), suggesting exaptive evolution of a zygnematophyceaen algae to glacial environments (Figure 2a, Supplementary Data 7, 9 & 13, Supplementary Information 5).

**Figure 2.**
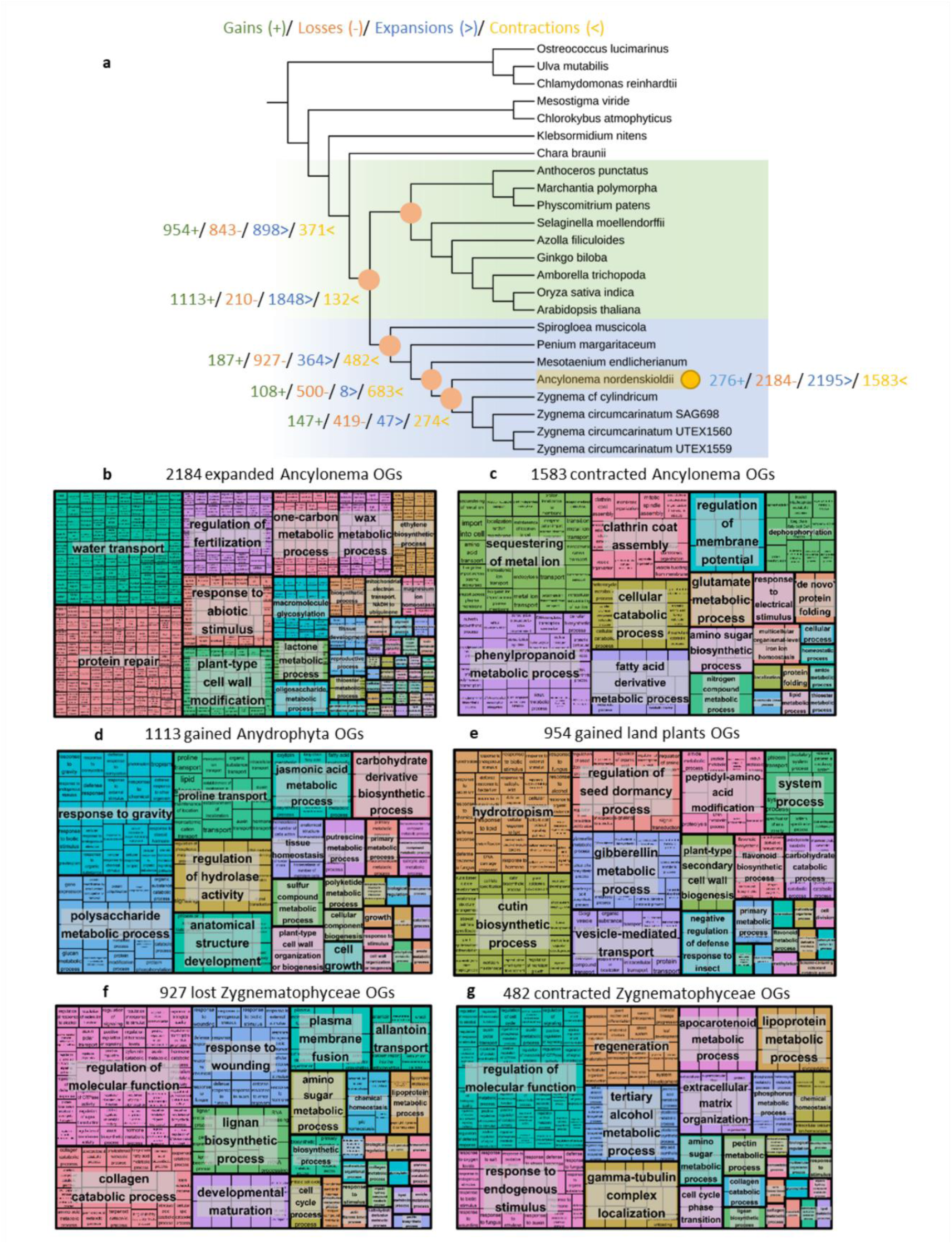
Comparative genomics of anydrophyte evolution. **A:** Gene family gain, loss, expansion and contraction across anydrophyte evolution. Key nodes are indicated on the tree. **B-G:** Gene Ontology analysis of **B:** expanded Ancylonema OGs, **C:** contracted Ancylonema OGs, **D:** gained Anydrophyta OGs, **E:** gained land plant OGs, **F:** lost Zygnematophyceae OGs & **G:** contracted Zygnematophyceae OGs.

**Figure 3.**
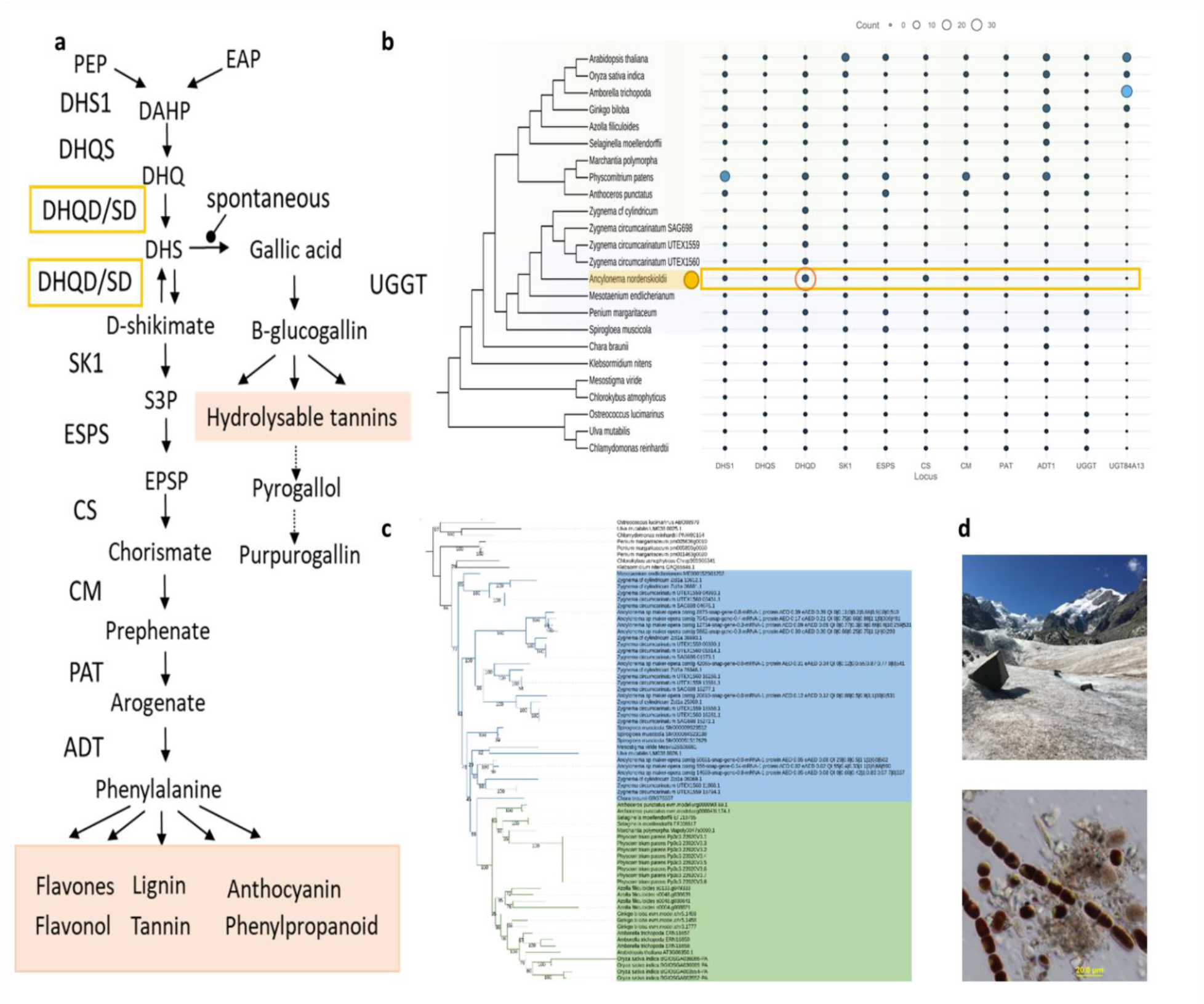
Comparative genomics of purpurogallin biosynthesis. **A,** Current understanding of the purpurogallin biosynthetic pathway. Orange boxes indicate secondary metabolites. **B,** Count of purpurogallin biosynthesis genes. The tree is based on phylogenetic analysis in Supplementary Information 4. The green background denotes land plants whilst blue background denotes Zygnematophyceae. Orange box highlights gene count for *Ancylonema nordenskiöldii.* Yellow circle highlights gene expansion of dehydroquinate dehydratase/shikimate dehydrogenase (DHQD/SD) in *Ancylonema nordenskiöldii.* **C,** Phylogenetic analysis of dehydroquinate dehydratase/shikimate dehydrogenase gene family. **D,** Illustration of pigmentation in glacier algae. The bottom image shows heavily pigmented glacier algae assemblages. The top image demonstrates surface ice environments of Morteratsch glacier with dark zone dominated by glacier algal blooms.

Our analysis also identified extensive gene loss and gene family contraction during the evolution of glacier algae (Figure 2). Gene losses and gene family contractions outweighed gains and expansions (Figure 2), suggesting extensive gene turnover with the evolution of glacier algae. Gene ontology analysis of 1583 contracted gene families revealed a reduction in functions associated with phenylpropanoid metabolism, clathrin coat assembly and cellular catabolic process (Figure 2c, Supplementary Information 10 & 14, Supplementary Information 5). GO terms associated with the 2195 lost gene groups, identified with Count^36^, mirrored the contracted gene families of *A. nordenskiöldii*. These lost genes were associated with phenol-containing compound metabolism, clathrin coat assembly and cellular catabolic assembly (e.g. cell wall organisation) as well as regulation of development (e.g. cell-cell signaling; Figure 2a, Supplementary Information 11 & 15, Supplementary Information 5). Together, these suggest a loss of intracellular trafficking, via clathrin coat assembly, and a reduction in chemical degradation pathways.

Despite the conspicuous phenolic pigmentation of *A. nordenskiöldii*, a known physiological adaptation to life in surface ice^21,37^, genes involved in the biosynthesis of many phenylpropanoids and phenolic compounds were lost in this species (Supplementary Information 5). Comparative genomic and phylogenetic analysis instead suggested that lineage-specific gene diversification of a particular pathway underpinned the novel screening pigmentation of *A. nordenskiöldii* (Figure 3). In glacier algae, purpurogallin pigments absorb ultraviolet and visible light, providing photoprotection against high levels of UV irradiance associated with life in supraglacial surface ice^37^. Purpurogallin accumulation has also been provisionally associated with protection against low temperatures, protection of the photosynthetic machinery and tolerance to desiccation^24^. Whilst genes involved in the purpurogallin biosynthetic pathway were found across the green plants, one component, dehydroquinate dehydratase/shikimate dehydrogenase (DHQD/SD), was present at a high copy number (9) in *A. nordenskiöldii*; greater than in any other green plant species (Figure 3, Supplementary Data 16). DHQD/SD catalyses the dehydration of dehydroquinate (DHQ) to dehydroshikimate (DHS) and the reduction of dehydroshikimate (DHS) to shikimate^38,39^. Phylogenetic analysis demonstrated that the expansion of this gene family was specific to *A. nordenskiöldii* (Figure 3c). After the synthesis of dehydroshikimate (DHS), gallic acid is spontaneously synthesised, eventually leading to the production of purpurogallin. Through gene duplication followed by neofunctionalisation, glacier algae are thus able to synthesise this novel pigment, facilitating their dominance of surface ice environments^18,21,24^. The high production of purpurogallin within glacier algal cells, i.e. to 11-times the cellular content of chlorophyll-*a*^21^, may have made other phenolic and phenylpropanoid-based compounds functionally redundant, explaining the loss and contraction of genes important for secondary metabolite biosynthesis highlighted in our comparative genomic analysis (Figure 2c).

Aside from purpurogallin, comparative genomic analysis did not find any other *A. nordenskiöldii* specific gene radiations in relation to light screening, including UV-A or UV-B stress (Supplementary Data 17), supporting previous assertions that purpurogallin provides the bulk of photo-protection through photo-quenching within glacier algal cells, whilst chloroplasts retain typically light-adapted for green algae^21^. This is further supported by genetic analysis of photosystem I and II here, showing no significant patterns of expansion or contraction in genes in the photosynthetic machinery (Supplementary Data 18).

Our analysis of ice-binding proteins suggested that cold adaptations of *A. nordenskiöldii* represent unique adaptation of glacier algae, rather than an ancient exaptation derived from Anydrophyta (Supplementary Data 19 & 20). Reciprocal BLAST of Uniprot ice-binding proteins from across the tree of life (e.g. fungi, bacteria) revealed 1194 OGs with hits against *A. nordenskiöldii* (Supplementary Data 19 & 20). Many of these were distributed across the green plant phylogeny, suggesting that more investigation into gene responses to freezing is required. These included a protein kinase superfamily, ATP-binding cassette protein family and heat shock protein family. Only 35 of these 1194 OGs were novel to Anydrophyta, indicating a relatively small evolutionary response to cold stress. Additionally, *A. nordenskiöldii* had the highest copy number amongst all green plants for some gene families, including a lipase, ATP-dependent Clp protease and Cytochrome C Oxidase.

Further analysis of land plant cold tolerance pathways did not find any gene family expansion in *A. nordenskiöldii*, indicating that its cryospheric adaptation likely did not derive from a mechanism common to embryophytes (Supplementary Information 6, Supplementary Data 21). Furthermore, this analysis indicated that a core element of land plant cold tolerance emerged after the split with Zygnematophyceae, arguing against the hypothesis^23,24^ that cold adaptations were exapted from a streptophyte algal ancestor. Cold stress signalling in land plants is coordinated by the CBF(C-REPEAT BINDING FACTOR)-COR(COLD REGULATED) signalling pathway^40^. Comparative genomics revealed that the majority of this pathway predated the origin of Anydrophyta (Supplementary Data 21). However, ICE (INDUCER OF CBF EXPRESSION), a key regulator of CBFs, emerged in the ancestor of land plants. ICE, also known as SCRM (SCREAM), is involved in stomatal development^41^, and may have evolved in land plants as an adaptation to both cold and drought stresses. Due to their importance in abiotic stress responses, the evolution of the biosynthesis and signalling of phytohormones was also investigated. Similar to previous analyses^12,28,42,43^, we demonstrated the prevalence of these phytohormone genes, which are mostly estimated to originate before the transition of plants onto land (Supplementary Data 22, Supplementary Information 7). CBF/DREBs (DEHYDRATION RESPONSIVE ELEMENT BINDING) are transcription factors that bind to DRE/CRT cis-acting elements in response to abiotic stress (e.g. drought, low temperatures). As such, the evolution of DREBs and all other transcription factors was investigated (Supplementary Data 23, Supplementary Information 8), with all major families being found in the *A. nordenskiöldii* MAG, consistent with previous studies^44–46^.

### Timescale of glacier algal evolution

Timescale analysis based on 472 genes from the genomes of 24 green plant species suggested that *Ancylonema nordenskiöldii* split from its closest living algal relative 455-520 million years ago (Ma) during the early Phanerozoic, emphasizing the sparseness of genome-scale sampling in this important region of the streptophyte tree (Figure 4). While no long-term glaciations occurred during this period, this estimate is predated and postdated by a number of glacial episodes (e.g. the mid Ediacaran Gaskiers, Late Ordovician and the late Carboniferous glaciations)^47^, potential drivers of glacier algal evolution. Additionally, regardless of their exact origination point, this represents a long evolutionary time in which to assemble glacial adaptations.

**Figure 4.**
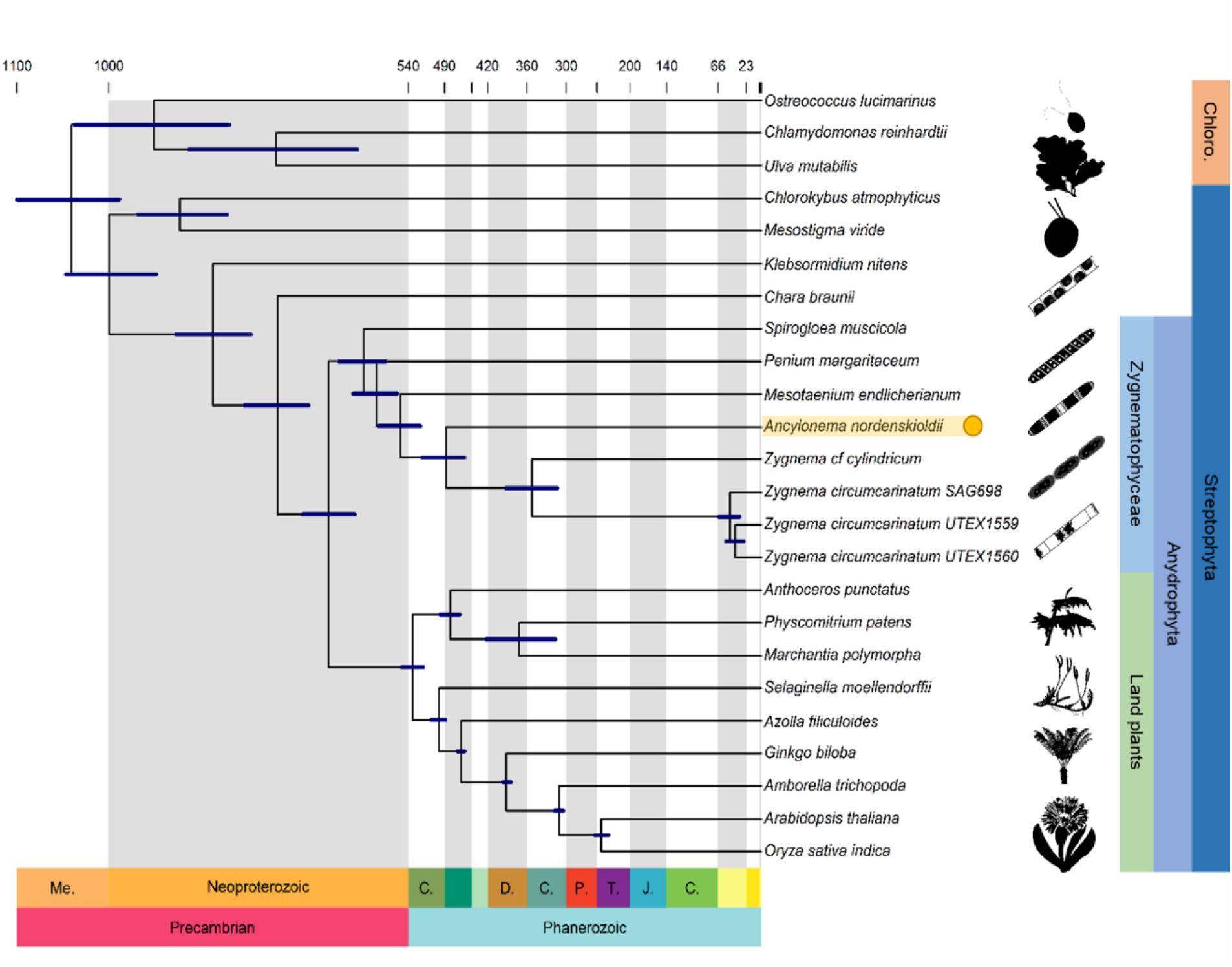
Timescale analysis of *Ancylonema nordenskiöldii.* Phylogenetic analysis of the placement of *Ancylonema* within the Anydrophyta, based on a concatenated alignment of 472 genes. *Ancylonema nordenskiöldii* highlighted in yellow. Taxonomic group are denoted on the side, with Chlorophytes abbreviated to Chloro.

Furthermore this timescale analysis, consistent with previous estimates^10,48,49^, suggested that Anydrophyta emerged 703-623 million years ago during the Cryogenian period (720 – 635 Ma), a period of dynamic environmental change characterized by two major global glaciations (Figure 4)^50,51^. While several studies produce an older estimate for crown Anydrophyta^11,52^, they require that key fossils (e.g. *Proterocladus antiquus, Bangiomorpha pubescens*) are assigned to derived phylogenetic positions that, in our view, are not justifiable based on the phenotypic evidence preserved (Supplementary Information 9). Our timescale analysis is potentially consistent with the hypothesis^23,24^ that Cryogenian glaciations drove the evolution of land plants. Aside from divergence time estimates, this hypothesis is based on the similarities of environmental constraints of the Cryogenian to those required for life on land (e.g. drought, high irradiance, freezing)^23,24^ and biomarker diversification^53,54^. These selective pressures would likely be reflected in the ancestral genomes of land plants and Zygnematophyceae. However, the above comparative genomic and phylogenetic analyses have suggested that adaptations to life in ice is lineage-specific to *Ancylonema*. As such, further comparative genomics (Supplementary Data 6, 7, 24-47) was used to investigate patterns and functions of genome evolution in Anydrophyta, land plants and Zygnematophyceae, and potential exaptive Cryogenian evolution.

### Genome and gene family evolution

Analysis of gene family evolution highlighted increasing genome complexity of Anydrophyta and land plants, contrasting with large-scale genome reduction in the Zygnematophyceae (Figure 2, Supplementary Data 24, 25, 28 & 29). The origin of Anydrophyta was accompanied by a peak of gene novelty (1113 gained & 1848 expanded OGs) and low gene loss (210 lost & 132 contracted OGs). These gained genes could derive from *de novo* formation from non-coding sequences, shuffling from a combination of existing domains or through horizontal gene transfer as recently identified in land plants^55–57^. Within the first-splitting lineages of Anydrophyta, the land plant ancestor exhibited another peak of gene novelty (954 gained & 898 expanded OGs). In contrast, within the ancestor of Zygnematophyceae gene loss and contraction were both high (927 lost & 482 contracted OGs; Supplementary Data 34 & 35). Further reductive evolution was also prevalent within the Zygnematophyceae, in the ancestor of Zygnematales (500 lost & 683 contracted OGs) and the common ancestor of *Ancylonema nordenskiöldii* and *Zygnema* species (419 lost & 274 contracted OGs; Figure 2a, Supplementary Data 6 & 7). These divergent genomic trajectories of Zygnematophyceae and land plants mirror their divergent morphological trajectories previously identified^25,26^ and suggests that genome streamlining may explain the reductive morphological evolution of Zygnematophyceae, resulting in the emergence of unicellular and filamentous algae from a multicellular Anydrophyta ancestor. Similar patterns of genome evolution have been observed in bryophytes, highlighting the importance of gene loss in plant evolution^48,58^.

Analysis of gene ontology (GO) terms and Pfam domains documented how ancestral anydrophytes and land plants evolved increasingly complex anatomies to establish in terrestrial environments (Figure 2d-e, Supplementary Data 36 & 40). Gained OGs in Anydrophyta were associated with responses to gravity, polysaccharide metabolic process (e.g. xyloglucan metabolism) and anatomical structure development (e.g. cell differentiation; Figure 2d, Supplementary Information 5). In land plants, gained OGs were associated with hydrotropism, cutin biosynthesis (e.g. meristem development), plant-type secondary cell wall biogenesis and system process (e.g. vascular and phloem transport; Figure 2e, Supplementary Information 5). It is likely that the evolution of xyloglucan metabolism and cutin biosynthesis led to the development of complex cell walls for protection against external stressors^59^. Mechanisms of detecting water, hydrotropism, as well as internal transport of water through xylem-like tissue also likely helped the first land plants increase in size and stature^45,60^.

Gene family expansions resulted in an increased number of stress response genes in the first Anydrophyta and in land plants (Figure 2a, Supplementary Data 37 & 41). Expanded OGs in Anydrophyta were linked to water transport, response to abiotic stimulus (e.g. responses to salt, osmotic stress, heat, cold) and plastid organisation (Supplementary Information 5). Land plant expanded OGs were associated with regulation of development, intercellular transport and response to abiotic stimulus (responses to salt, temperature, osmotic stress; Supplementary Information 5). Through gene family expansions, a greater repertoire of stress response pathways emerged in Anydrophyta and land plants, which potentially enabled them to tackle temperature, drought and osmotic stress of the Cryogenian through to the Cambrian. While our analysis found no clear evidence of the ancient cryosphere in driving anydrophyte diversification, the physical environment clearly poses selection pressures for plant evolution, the exact nature of which is yet to be determined.

Our GO analysis suggested that extensive loss and contraction of gene families in the first Zygnematophyceae enabled the reductive evolution of their simple morphology from a more complex ancestor (Figure 2, Supplementary Data 46 & 47). The OGs lost from Zygnematophyceae were associated with regulation of molecular function (e.g. regulation of cell communication), collagen catabolic process, lignan biosynthesis and plasma membrane fusion (e.g. extracellular matrix and structure organisation; Figure 2f, Supplementary Information 5). Contracted OGs in Zygnematophyceae were linked with regulation of molecular function (e.g. auxin polar transport), responses to endogenous stimuli (e.g. salt stress, hypoxia, biotic stimulus), regeneration (e.g. multicellular organism development) and extracellular matrix organisation (Figure 2g, Supplementary Information 5). As extant Zygnematophyceae algae are unicellular or filamentous^25^, the loss and contraction of gene families involved in extracellular matrix organisation provides a clear molecular signature of the evolution of their simple morphology. These losses include the synthesis of cell wall compounds, as well as the capacity to communicate between cells and organize multicellular structure.

## Conclusions

Here, we produced the first genome-scale data for glacier algae, which occupy a climatologically important niche^21^. Analysis incorporating green plant genomes revealed a divergent genomic trajectory of land plants and Zygnematophyceae from their common ancestor, which corresponds to morphological data. Contrasting patterns of genome evolution between closely related groups have been seen elsewhere in the tree of life^48,61–63^, highlighting the role of gene gains as well as losses in driving phenotypic evolution. Our gene ontology analysis revealed that gene gains in Anydrophyta and land plants were associated with anatomical structural development while in Zygnematophyceae, gene losses were linked to multicellular development. This suggests that modern Zygnematophyceae evolved by reductive evolution from ancestors with larger gene repertoires, a pattern also reported in bryophytes^48^. OGs gained and expanded in Anydrophyta were not linked to ice stress tolerance, but rather specific adaptations to the cryosphere appeared to be a novelty of glacier algae. Adaptation to ice in *Ancylonema nordenskioldii* did not result from exaptation (that is, modification of an ancient gene repertoire). This was further elucidated by inferences that lineage-specific gene diversification likely led to their novel pigmentation that protects them from environmental extremes associated with glaciers. Thus, the comparatively recent ice adaptation of *Ancylonema* allowed us to reject the hypothesis that they are a vestige of a Cryogenian anydrophyte. Despite this, many expanded gene groups in Anydrophyta and land plants were linked to abiotic stress responses, such as drought, osmotic and temperature stress. This adds to the growing body of work demonstrating the stress tolerance capabilities of the common ancestor of land plants and Zygnematophyceae^13,15,16,64,65^, eventually leading to the establishment of plants on land. Indeed, this bears out the expectation that the freshwater algal relatives of embryophytes were confronted by much the same challenges as their land plant relatives, and the adaptive solutions that they established preadapted stem-embryophytes for life on land^66^.

## Methods

### Study site and sample collection

Communities of ice-inhabiting glacier algae were sampled from the surface ice of Morteratsch glacier, Switzerland, in August 2020. Surface ice was collected using a pre-sterilized sampling ice-saw directly into sterile Whirl-Pak bags and maintained frozen at -20°C during transport to the University of Bristol, UK, whereby samples were held at -80°C prior to genomic DNA extraction as below.

### Library construction and sequencing

For short-read sequencing, genomic DNA was extracted using the DNeasy PowerSoil Pro kit (Qiagen, USA) and assessed with Qubit (ThermoFisher Scientific, USA), BioAnalyser (Agilent, USA) and Femto Pulse (Agilent, USA). Samples were purified through bead-based clean up and PCR-free libraries were prepared by Earlham Institute. Short-read libraries were sequenced with Illumina NovaSeq 6000 system in paired-end mode. In total, 369.4 Gb raw data were generated, with 333.7 Gb remaining after filtering by Trimmomatic^67^.

For long-read sequencing, samples were ground in liquid nitrogen for 15 minutes and genomic DNA was extracted following a previously described protocol^68^ and assessed with Qubit (ThermoFisher Scientific, USA), BioAnalyser (Agilent, USA) and Femto Pulse (Agilent, USA). The NERC Environmental Omics Facility sequenced long-read libraries with PacBio, producing 22.2 Gb raw data.

### Genome assembly

Short (Illumina) and long read (PacBio) datasets were assembled with hybrid assembler OPERA-MS^69^. A minimum read length of 3000 bp was specified. As the sequenced samples represented communities of organisms, a metagenomic approach was used to extract streptophyte algal data. Eukrep^70^ was used to classify eukaryote contigs. Within these eukaryotic reads, Kraken2^71^ was used to identify 9979 streptophyte contigs. Multiple methods were then used for genome binning, including Maxbin^72^, Metabat2^73^, BinSanity^74^ and Concoct^75^. These were evaluated with the BUSCO Eukaryota dataset^29^ to identify Metabat2 as the best binning tools (Supplementary Information 1). Genome qualities was also confirmed with BUSCO Viridiplantae (Supplementary Data 5).

### RNA sequencing and assembly

RNA for Illumina RNA-Seq was extracted with the RNEasy PowerSoil Total RNA kit (Qiagen, USA) following the manufacturer’s protocol. The quality of RNA was assessed using an Agilent BioAnalyzer (Agilent, USA). The Bristol Genomic Centre sequenced RNA-Seq libraries with Illumina HiSeq 2500 system in paired-end mode. RNA sequencing data were mapped against our streptophyte algal MAG with HISAT2^76^, using default parameters. Mapped reads were then assembled with Trinity^77^, with default parameters.

### Gene prediction

Repetitive elements were identified with RepeatModeler^78^ and masked from the original genome assemblies with RepeatMasker^79^. The MAKER-P^80^ pipeline was used for gene annotation in two rounds, incorporating multiple annotation sources. Firstly, assembled RNA reads were mapped to the genome using TopHat2^81^. Homology-based gene prediction was completed with the protein coding genes of closely related species including *Mesotaenium endlicherianum*^27^, *Penium margaritaceum*^28^ and *Spirogloea muscicola*^27^. Gene models obtained from RNA-aided and homology-based pipelines were used to train the *de novo* prediction pipeline, SNAP^82^. A final MAKER-P run, combining all these sources, was used to annotate genes for the *Ancylonema nordenskioldii* MAG^80^. Genome completeness was assessed with BUSCO Eukaryota^29^. Functional annotation of protein coding genes was completed with InterproScan^83^.

### Distinguishing species of microalgae

Morphological analysis has identified two common species of glacier algae, unicellular *Ancylonema alaskanum* and filamentous *A. nordenskioldii*. Marker based analysis of rbcL and 18S genes demonstrated that these species of *Ancylonema* are very closely related^33^. To date, this is the only published genetic data for glacier algae. BLAST comparison of rbcL and 18S gene sequences supposedly from *Ancylonema alaskanum* and filamentous *A. nordenskioldii* data against the algal metagenome finds best hits against the same contig (Supplementary Information ?). The best hits identified our MAG as *A. nordenskiöldii* (Supplementary Data 1-4). Additionally, analysis was used to confirm a MAG deriving from a single species. GC content analysis, using SeqKit^84^, and kmer analysis, using Jellyfish^85^, were conducted. The function of duplicated BUSCO Viridiplantae genes were also assessed, using functional categories in OrthoDB^86^.

### Phylogenetic Analysis

We utilised the latest transcriptome and genome data to infer the evolutionary history of streptophytes. Specifically, these included Zygnematophyceae transcriptomes from the one thousand plant transcriptomes project (1KP) and genomes across the plant tree of life (Supplementary Information 3). A benchmark of 30% BUSCO genes missing was used to filter high quality data^29^. Two datasets were produced, the first constructed based on genome only data from across the green plants and the second constructed from Zygnematophyceae data from the 1KP project and complementary Anydrophyta genome data.

For both datasets, OrthoFinder (Version 2.3.7) was used to cluster protein coding genes into orthogroups^87^, using default settings (orthofinder -f data_folder). Single copy orthologs were identified using a previously described python script^88^, which removes paralogous genes from orthogroups. The script enables the user to specify a minimum taxonomic occupancy of each orthogroup, set at 70%.

Single copy orthologs were aligned using MAFFT^89^ using –auto parameter and trimmed with Trimal^90^ using the –automated1 parameter. Two complementary approaches were used to reconstruct phylogenies. In the first approach (concatenation), multiple sequence alignments were concatenated using Phyutility to create a supermatrix^91^. A bootstrapped maximum likelihood phylogeny was inferred using IQ-Tree^92^ using the Bayesian Information Criterion (BIC) to select best fitting substitution model and empirical profile mixture models (C10-C60). 1000 ultrafast bootstrap replicates were used. In the second approach (coalescent), individual bootstrapped maximum likelihood phylogenies were inferred using IQ-Tree as described above. These individual phylogenies were then summarised into a species tree using ASTRAL^93^.

### Species Divergence Time estimation

For molecular clock analysis, we utilized the aligned and trimmed 472 gene dataset from the green plant genome phylogeny as input into MCMCtree^94^. Node distributions using minimum and maximum constraints were specified, with full phylogenetic and age justifications listed in Supplementary Information 9. These calibrations derive from previous critical reviews of the fossil record^26,48^. To specify the prior distributions on node ages, all calibrated nodes were given a hard minimum age and a soft maximum age.

Initially, analyses were run without sequence data to obtain effective time priors, to ensure that the calibration densities and time priors were appropriate. The single copy orthogroups were divided into 4 partitions according to their evolutionary rate, based on total tree length in IQTree^92^ and grouped using k-means clustering in R^95^. A relaxed clock model was used (Uncorrelated; Independent Gamma Rates). Given the protein dataset, branch lengths were first estimated using codeml^94^. The tree topology was fixed based on the focal maximum likelihood analysis above and was analysed using the normal approximation method in MCMCtree^94^. After a burn-in of 10,000 generations, parameter values were saved every 20^th^ generation until 20,000 cycles were saved (400,000 generations total). Trees were plotted using MCMCTreeR^96^.

### Gene family evolution analyses

The gain, loss, expansion and contraction of gene families across the green plant genomes was analysed. Using the dataset containing green plant genomes, Count^36^ was used to analyse the gain and loss of genes with Dollo parsimony using the concatenation based phylogenetic tree produced above and the orthogroup count table from Orthofinder as input. CAFÉ^35^ was used to analyse gene family expansion and contraction, using the previously produced time-calibrated phylogeny (Figure 4) and the orthogroup count table from Orthofinder as input using default parameters.

Protein domains were also assessed with Interproscan^83^. The function of gained, lost, expanded and contracted gene families were visualized with REVIGO^98^ (using Gene Ontology terms from Interproscan).

### Evolution of key gene families

The occupancy of gene families involved in key biological processes were queried within the outputs of Orthofinder. Glacier algae are found across the cryosphere and tolerate extremes of temperature, desiccation and UV radiation. Thus, these focused on transcription factors, phytohormone biosynthesis and signaling, circadian rhythm, secondary metabolite biosynthesis and photosynthesis (Supplementary Data 16-19 & 21-23). Further inspection by gene tree analysis was completed in IQ-Tree using the same criteria as for the species tree analysis.

## Data availability

The *Ancylonema nordenskioldii* MAG, associated proteins, and genes used to infer species phylogenies were deposited in publicly available repositories (ENA: PRJEB67662 and Figshare: Project 183640).

## Author contributions

The work was devised by AMCB, TAW, PCJD, DAC & CJW. AMCB conducted the analyses and drafted the manuscript. All authors contributed to writing, reviewing and editing the finished manuscript.

## Acknowledgements

We wish to acknowledge funding from the Leverhulme Trust (RPG-2020-199 “iDAPT” project to CW, DC, PD, TW; RF-2022-167 to PD); the Natural Environment Research Council (NE/P013678/1 to PD; part of the Biosphere Evolution, Transitions and Resilience (BETR) programme cofunded by the Natural Science Foundation of China (NSFC); the John Templeton Foundation (62220 to PD, TW; the opinions expressed here do not necessarily reflect the views of the John Templeton Foundation); the Gordon and Betty Moore Foundation (GBMF9741 to PD, TW) and a University Research Fellowship to TW (URF\R\201024).

