## Supplementary Information for "Genome of the glacier alga *Ancylonema* and its insights into the evolution of streptophyte life on ice and land"

**Supplementary Information**
**Supplementary Information 1**

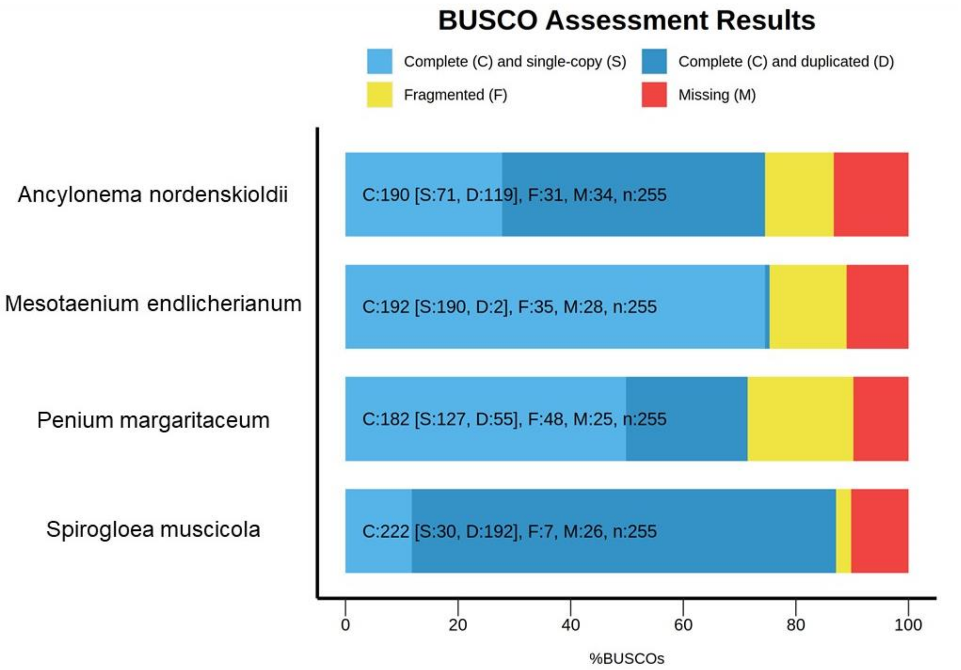


**Supplementary Figure 1.** BUSCO analysis of *Ancylonema nordenskioldii* MAG, as well as previously published Zygnematophyceae genomes[1,2].

**Supplementary Information 2**

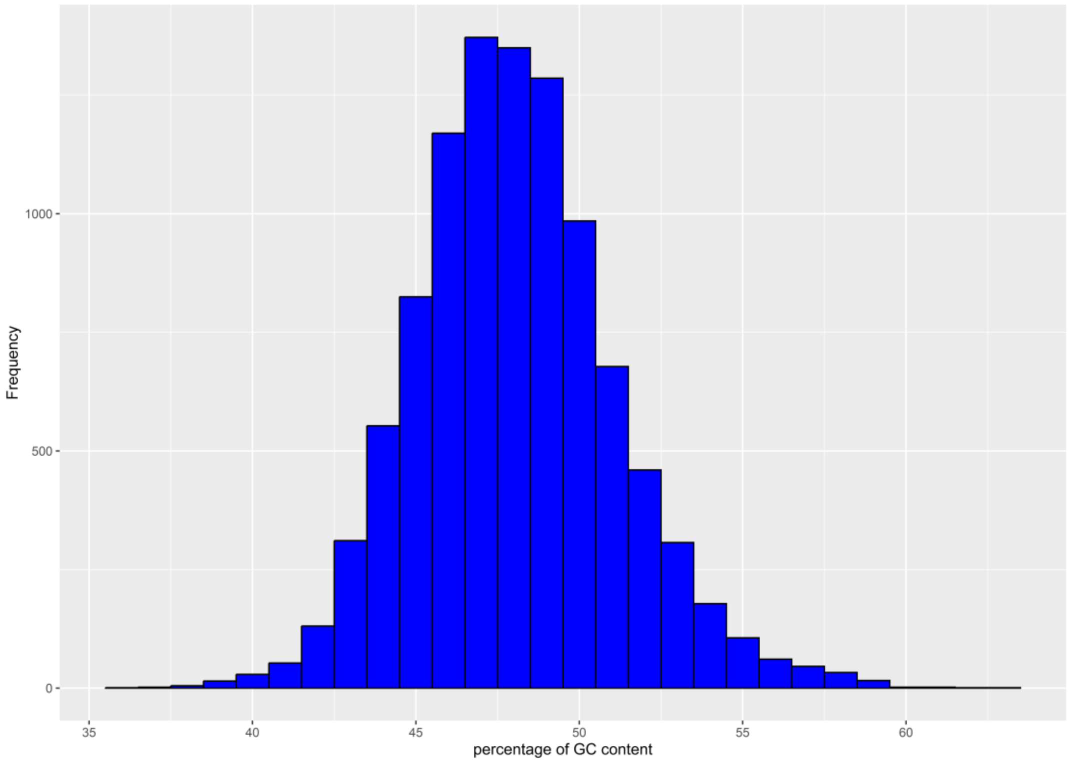


**Supplementary Figure 2.** Analysis of GC content for the contigs of *Ancylonema nordenskioldii* MAG.

**Supplementary Information 3**

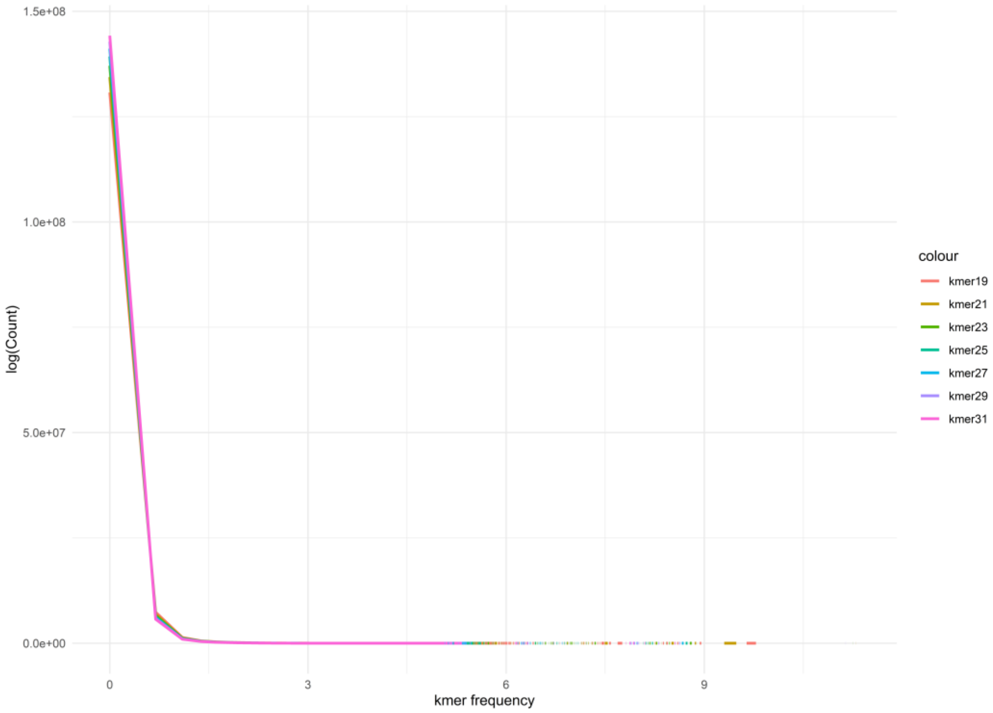


**Supplementary Figure 3.** Analysis of kmers for the contigs of *Ancylonema nordenskioldii* MAG

### Supplementary Information 4

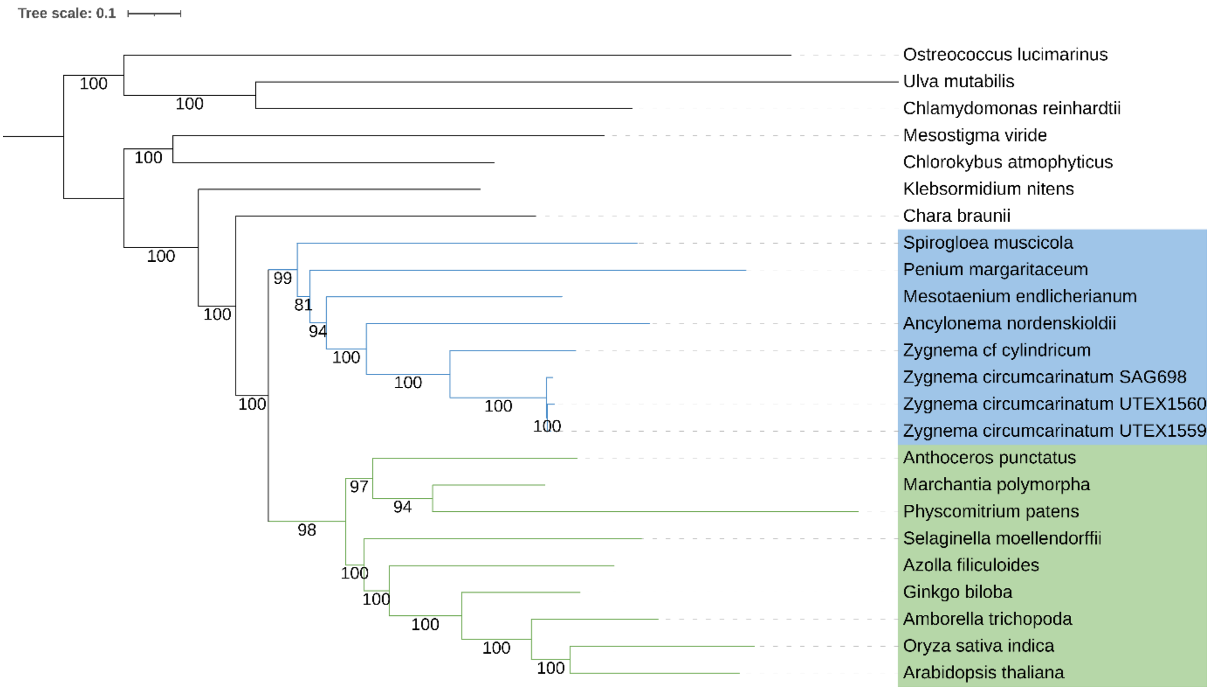


**Supplementary Figure 4.** Phylogenetic analysis of green plant genomes[1–18] based on 472 genes using a concatenation based approach. Numbers indicates bootstrap support values. Members of the Zygnematophyceae are shaded in blue, whilst land plants are shaded in green.

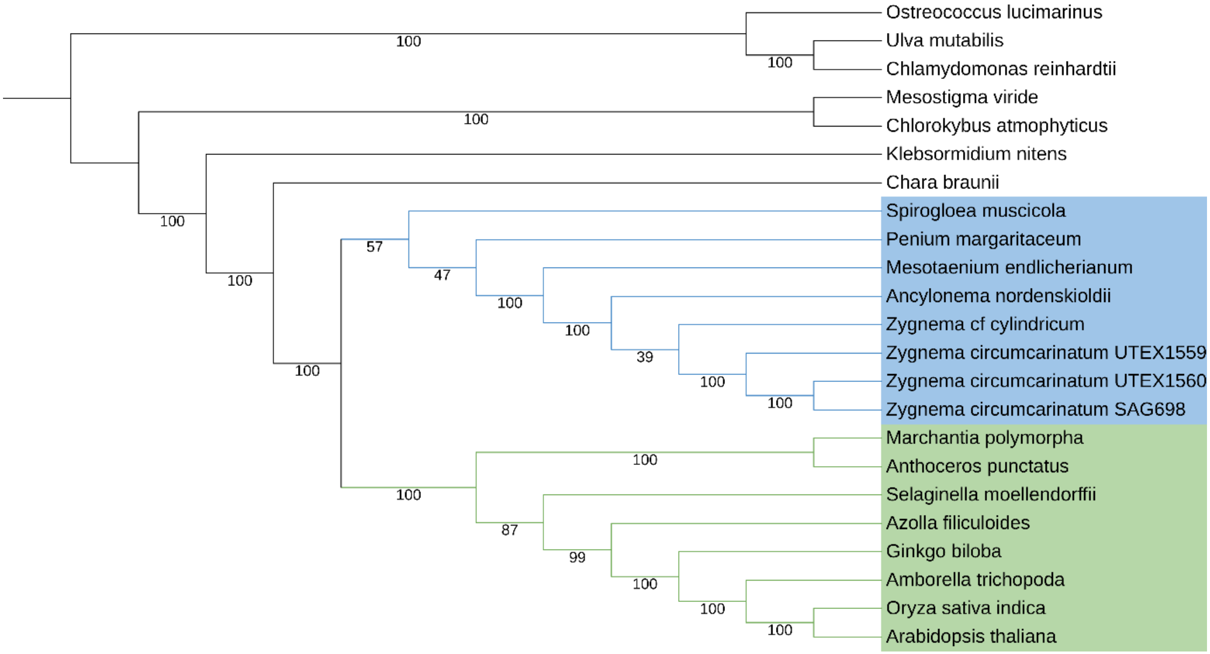


**Supplementary Figure 5.** Phylogenetic analysis of green plant genomes[1,2,11–18,3–10] based on 472 genes using a coalescence based approach (ASTRAL)[19]. Numbers indicates bootstrap support values. Members of the Zygnematophyceae are shaded in blue, whilst land plants are shaded in green.

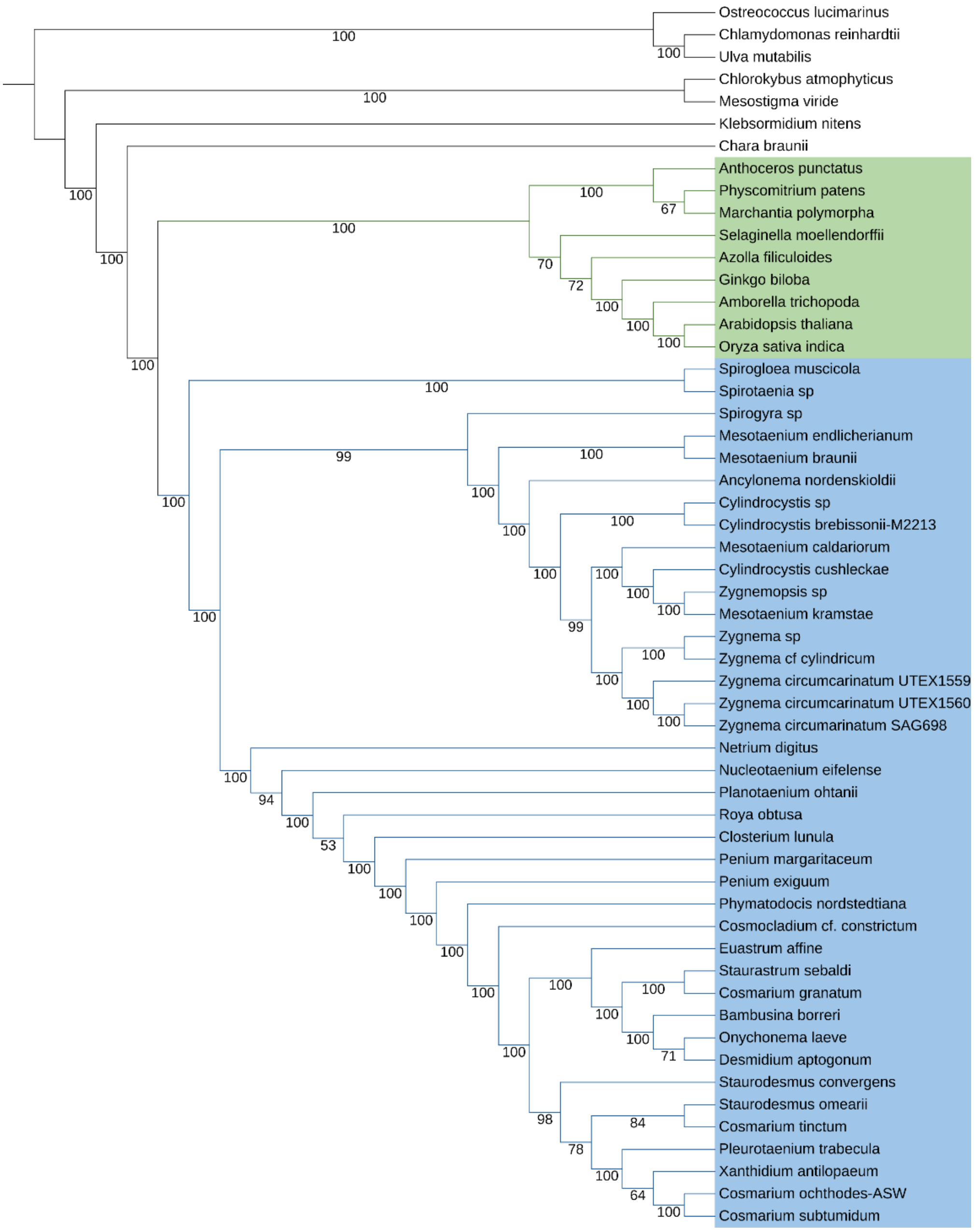


**Supplementary Figure 6.** Phylogenetic analysis of green plants incorporating data from the 1KP project[20] as well as genome data[1,2,11–18,3–10]. Analysis is based on 543 genes using a coalescence based approach (ASTRAL)[19]. Numbers indicates bootstrap support values. Members of the Zygnematophyceae are shaded in blue, whilst land plants are shaded in green.

**Supplementary Information 5 – REVIGO (Reduced visualisation of GO terms) plots**

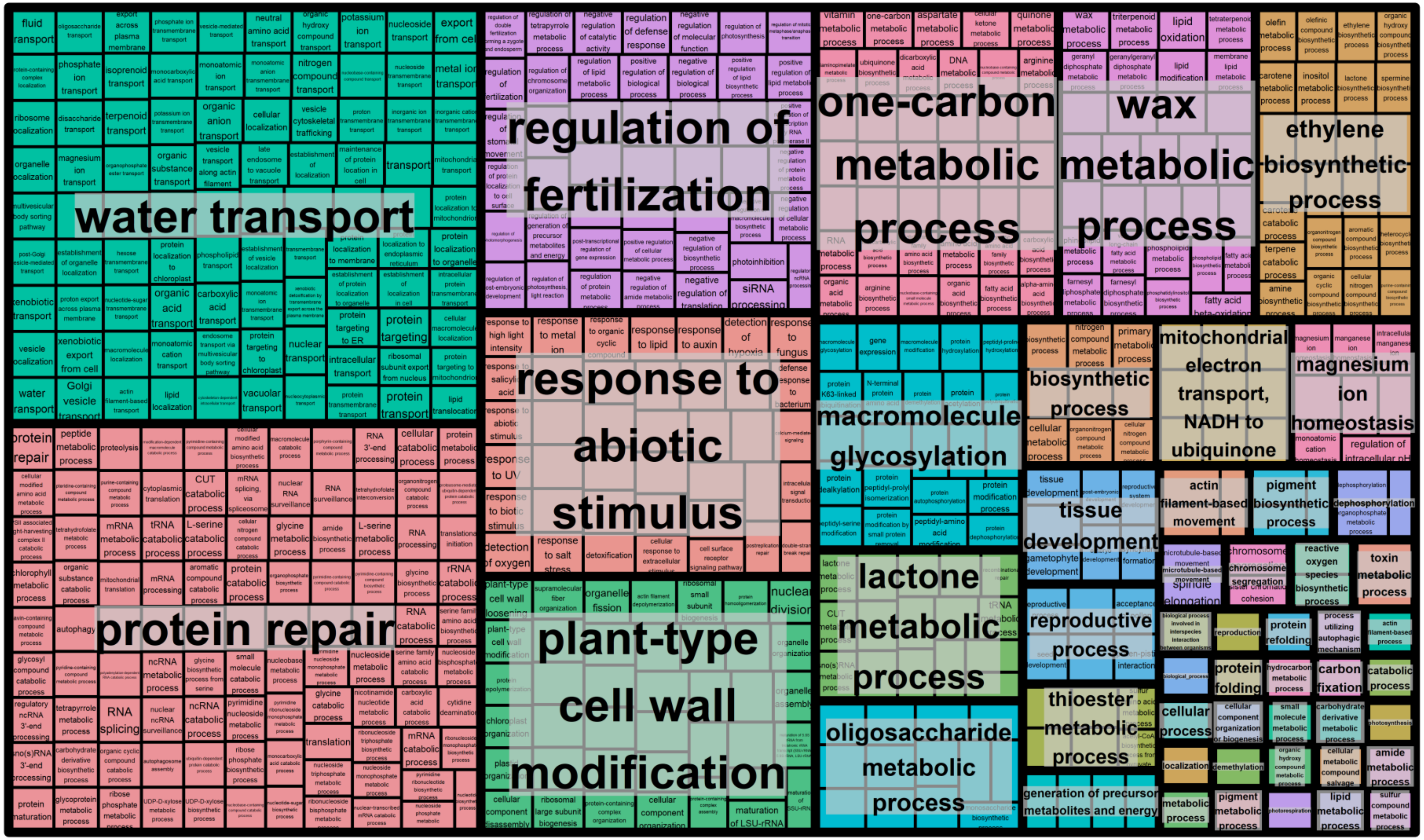


**Supplementary Figure 7.** REVIGO plot for expanded gene families in *Ancylonema nordenskioldii*

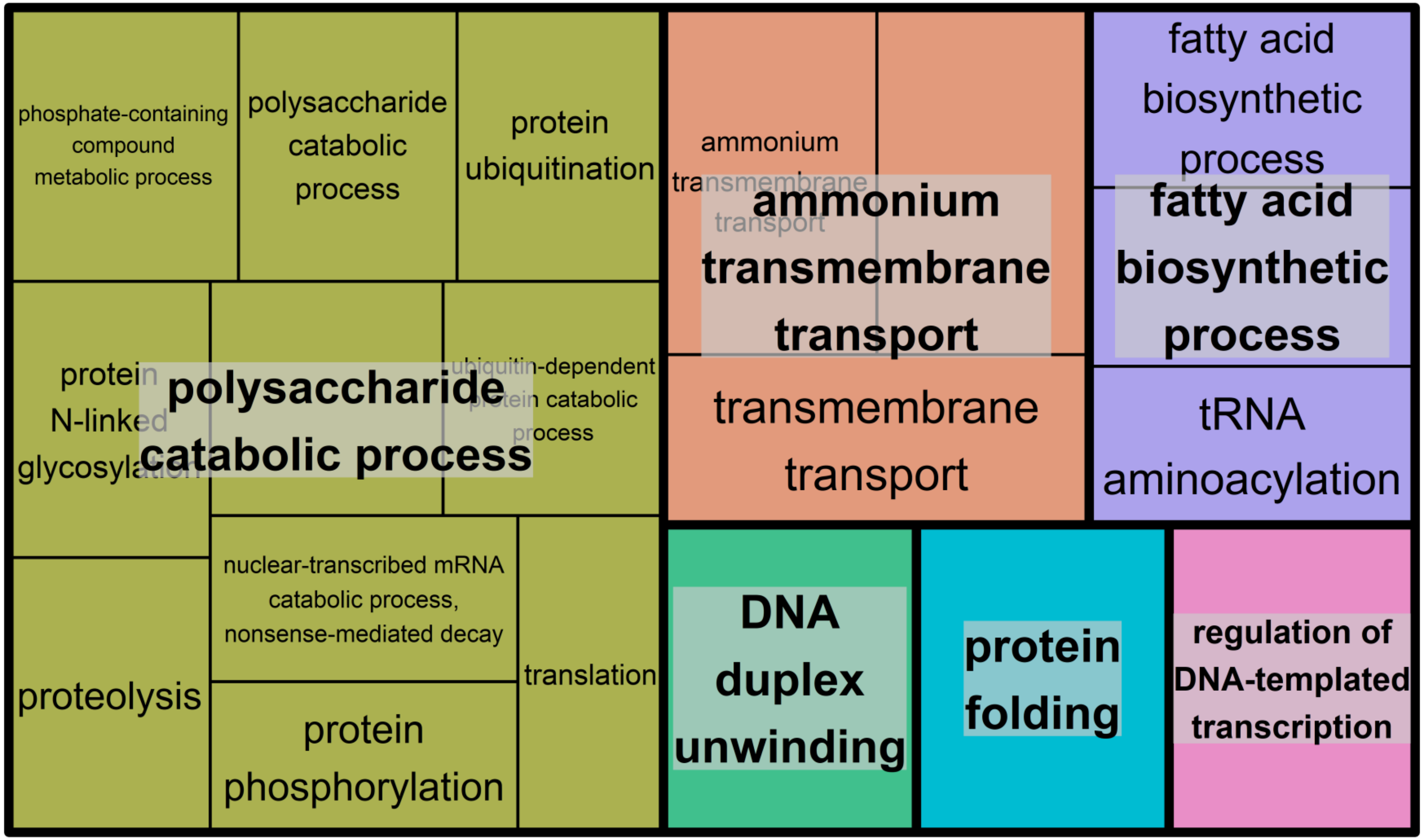


**Supplementary Figure 8.** REVIGO plot for gained gene families in *Ancylonema nordenskioldii*

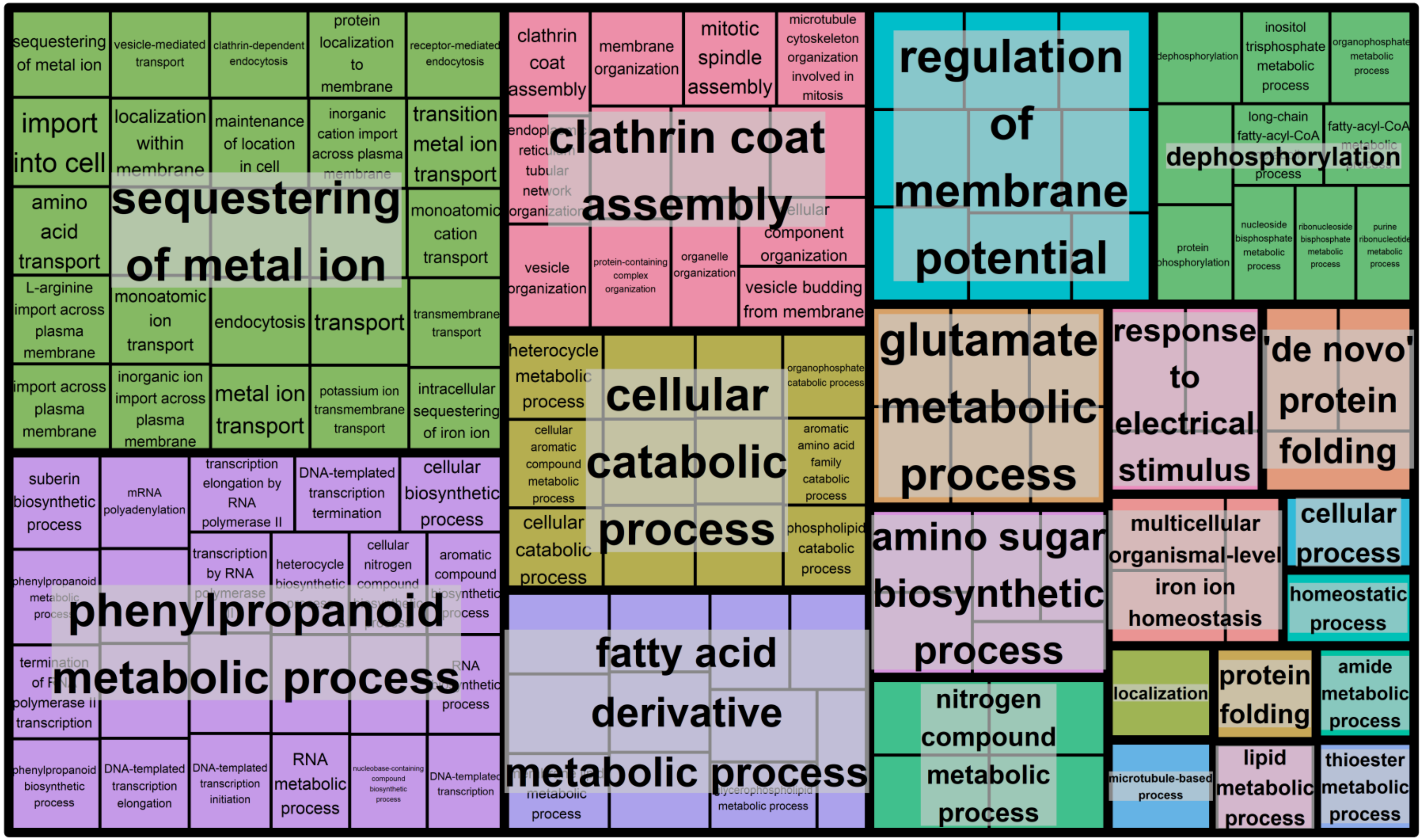


**Supplementary Figure 9.** REVIGO plot for contracted gene families in *Ancylonema nordenskioldii*

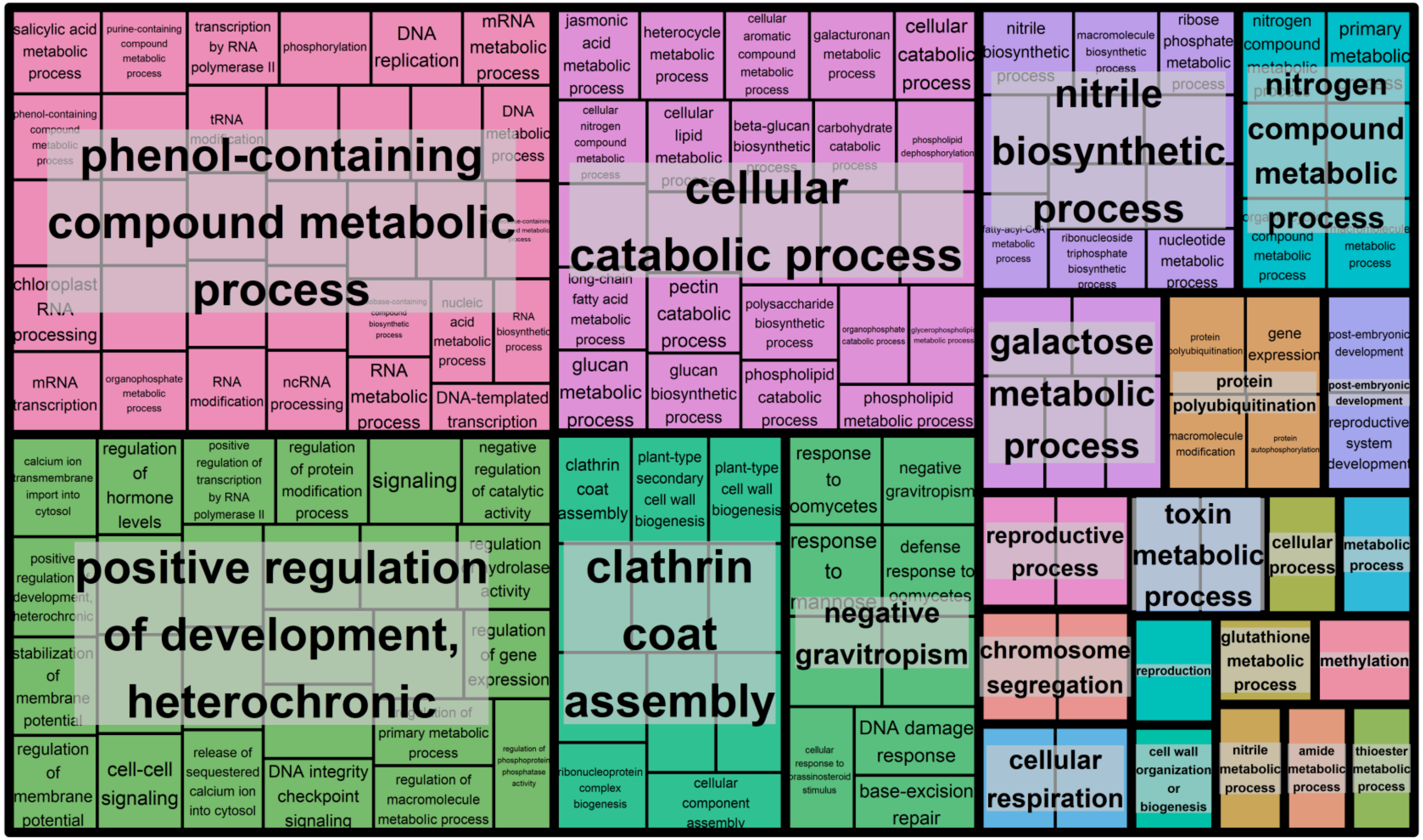


**Supplementary Figure 10.** REVIGO plot for lost gene families in *Ancylonema nordenskioldii*

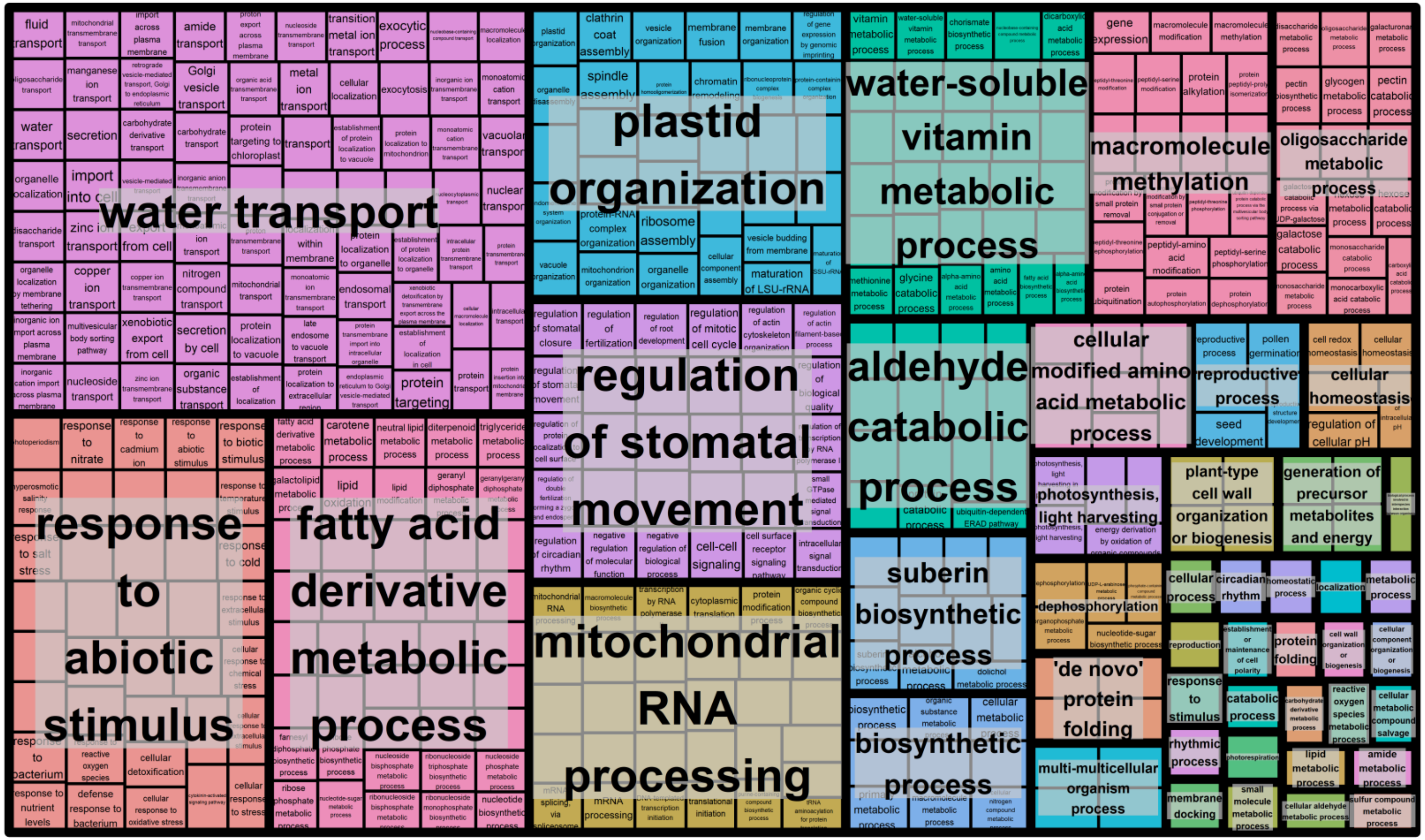


**Supplementary Figure 11.** REVIGO plot for expanded gene families in Anydrophyta

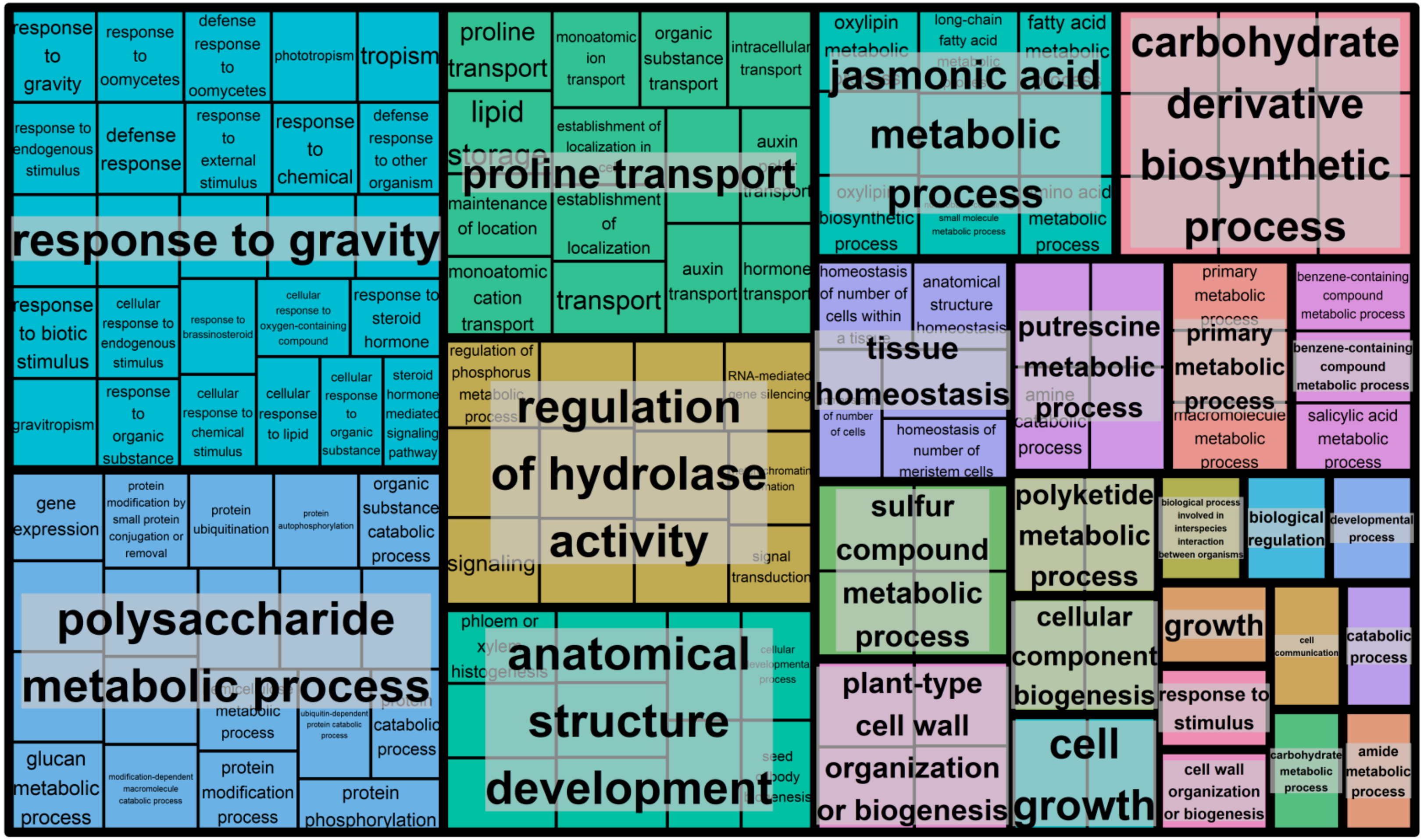


**Supplementary Figure 12.** REVIGO plot for gained gene families in Anydrophyta

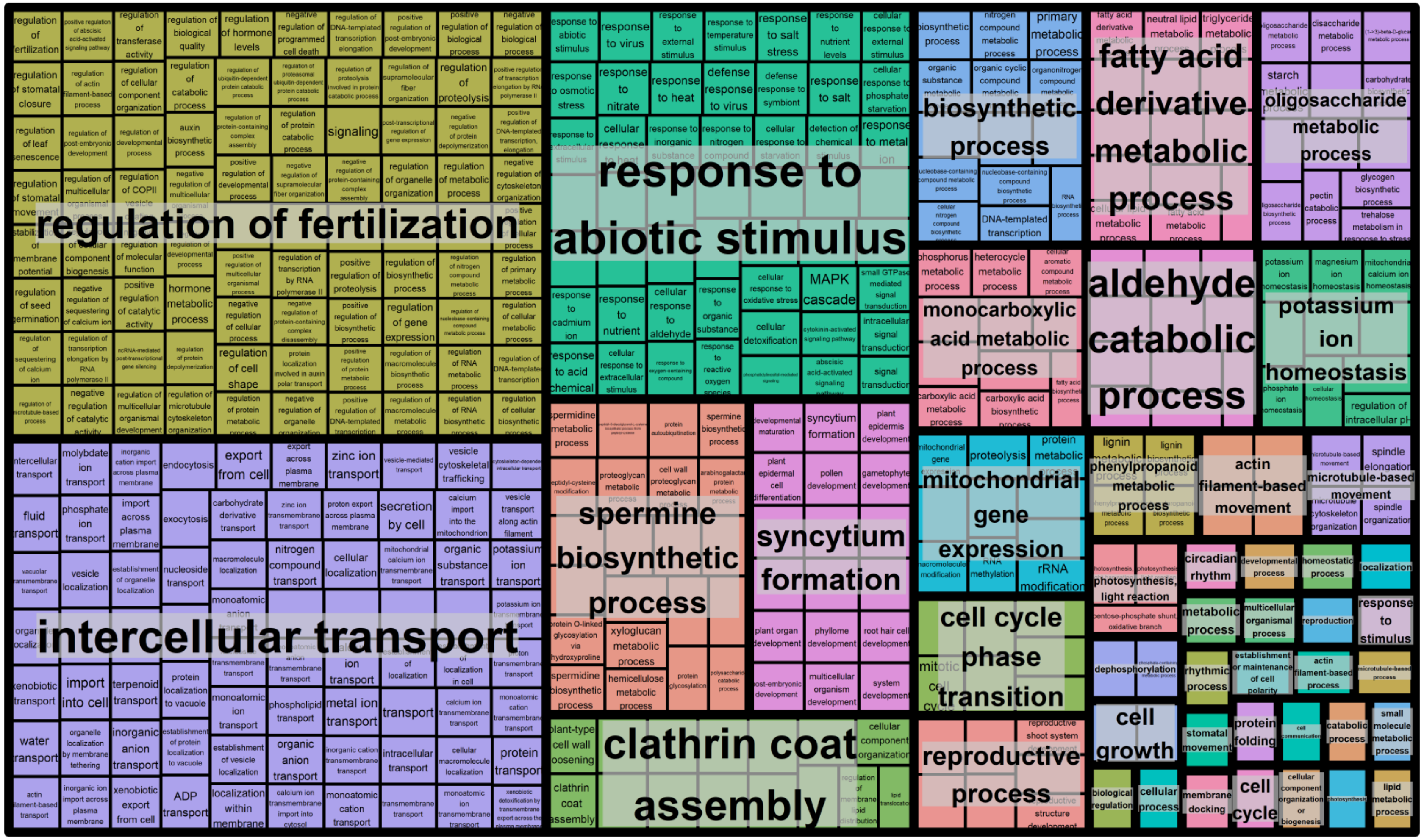


**Supplementary Figure 13.** REVIGO plot for expanded gene families in land plants

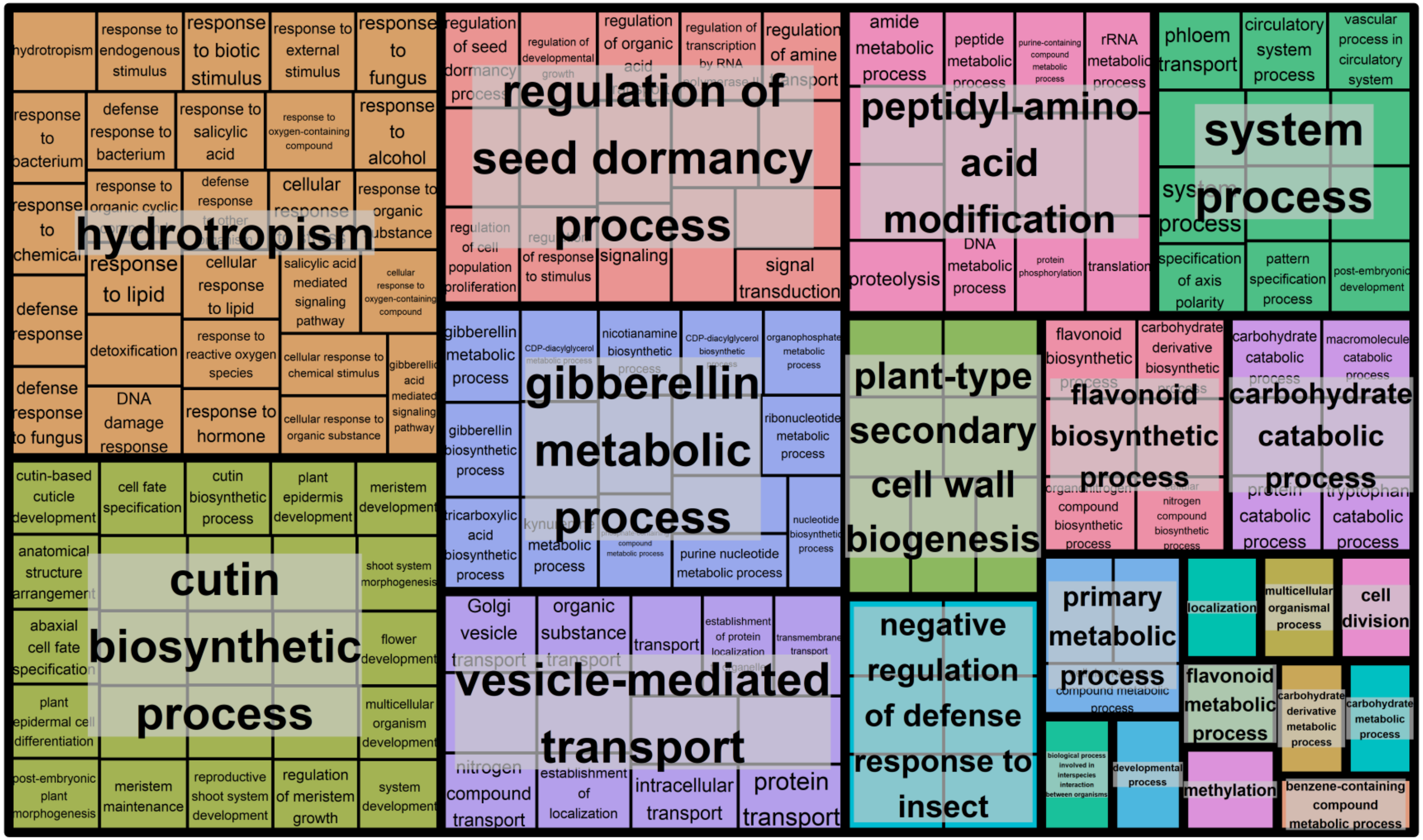


**Supplementary Figure 14.** REVIGO plot for gained gene families in land plants

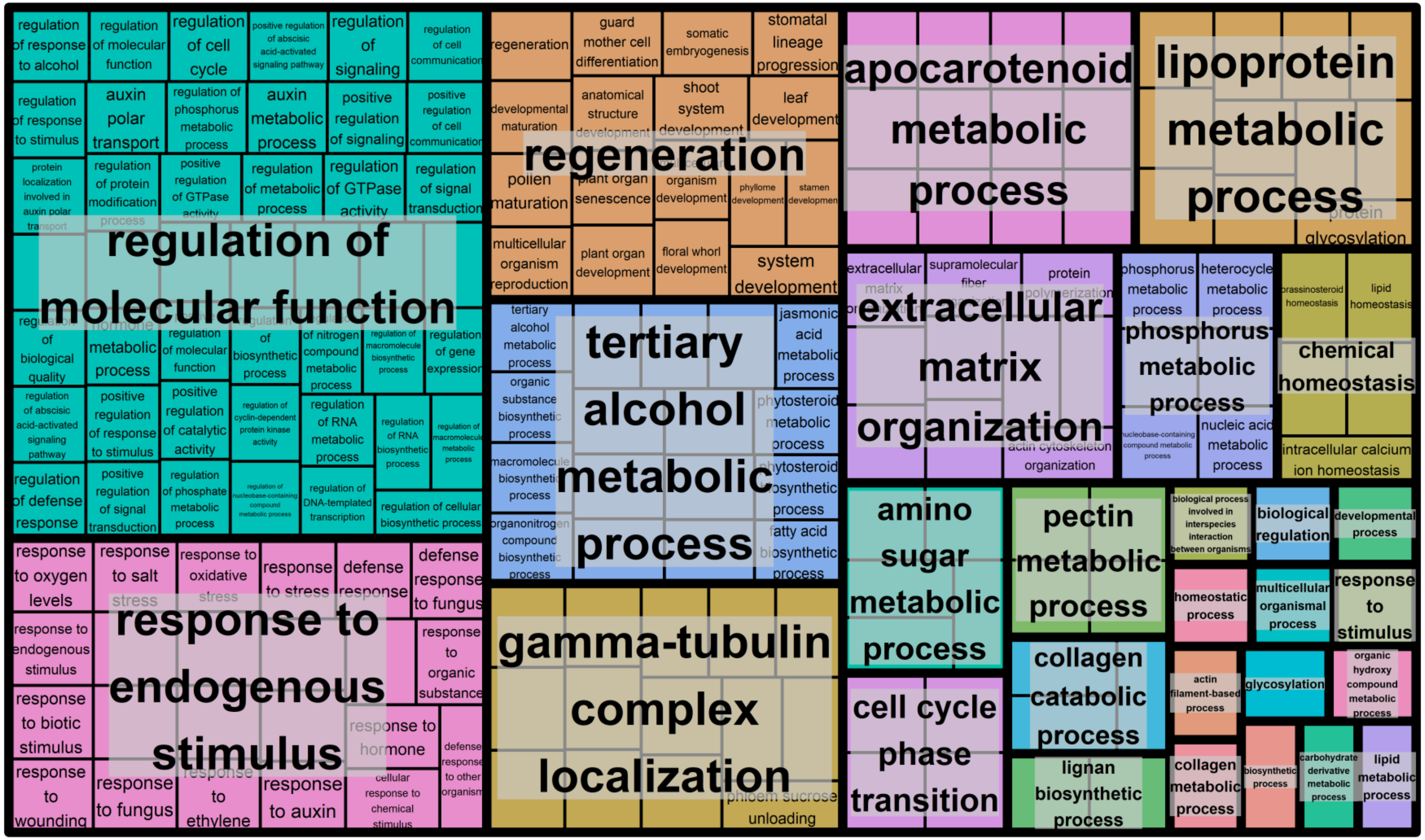


**Supplementary Figure 15.** REVIGO plot for contracted gene families in Zygnematophyceae

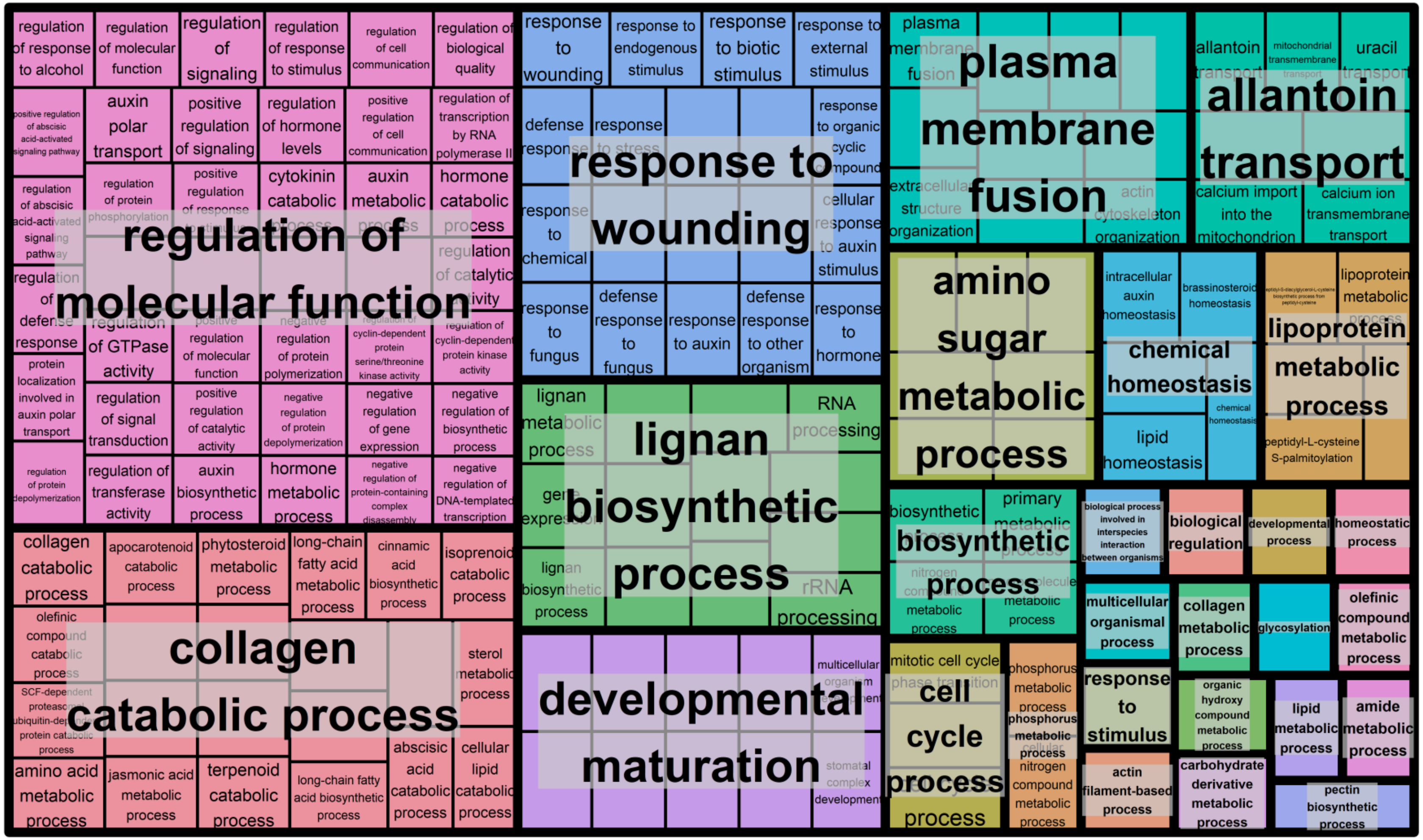


**Supplementary Figure 16.** REVIGO plot for lost gene families in Zygnematophyceae

**Supplementary Information 6**

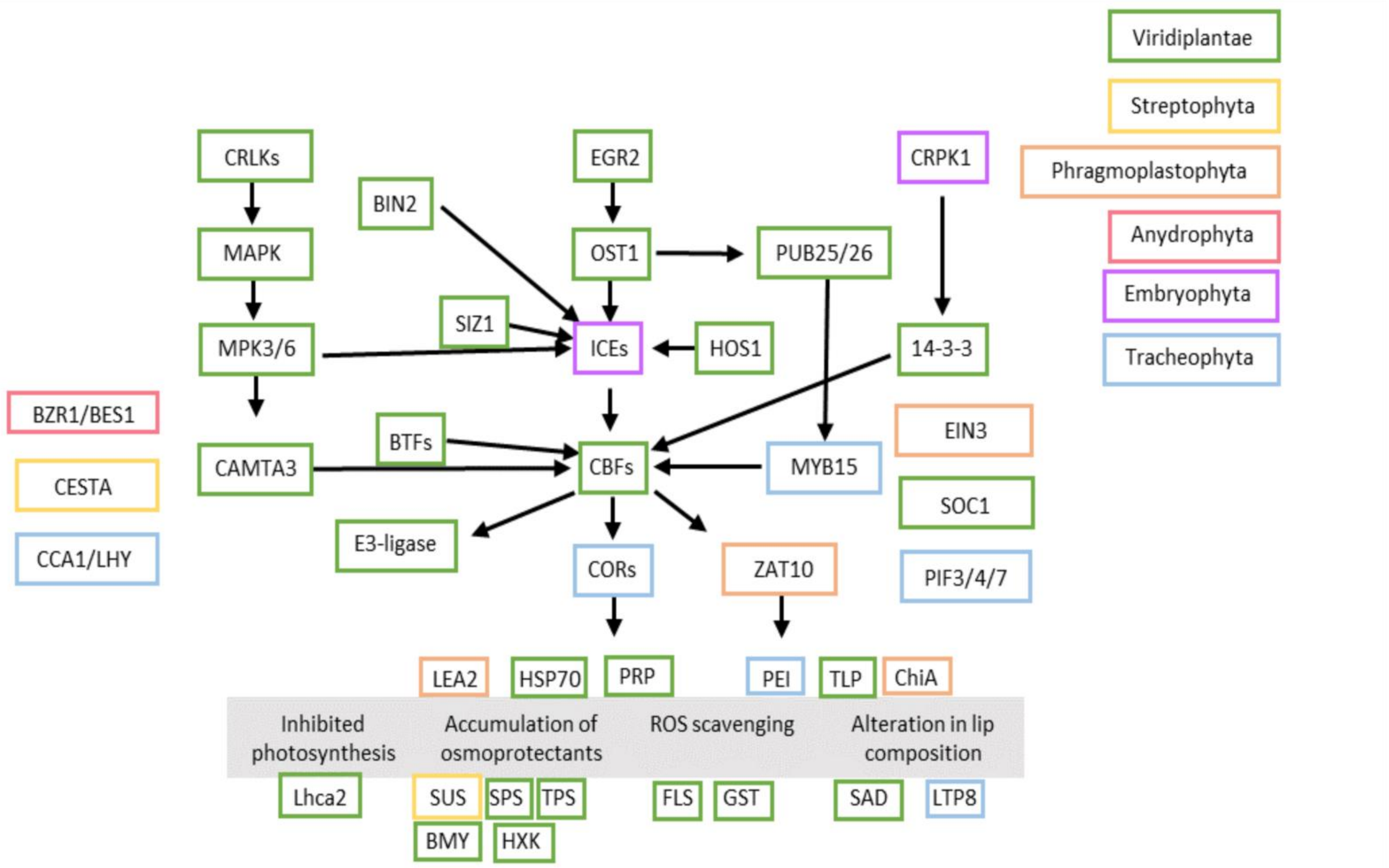


**Supplementary Figure 17** The evolution of land plant cold stress signalling pathways. Each box refers to a gene and is colour coded by its phylogenetic origin, highlighted in the key in the top right.

**Supplementary Information 7**

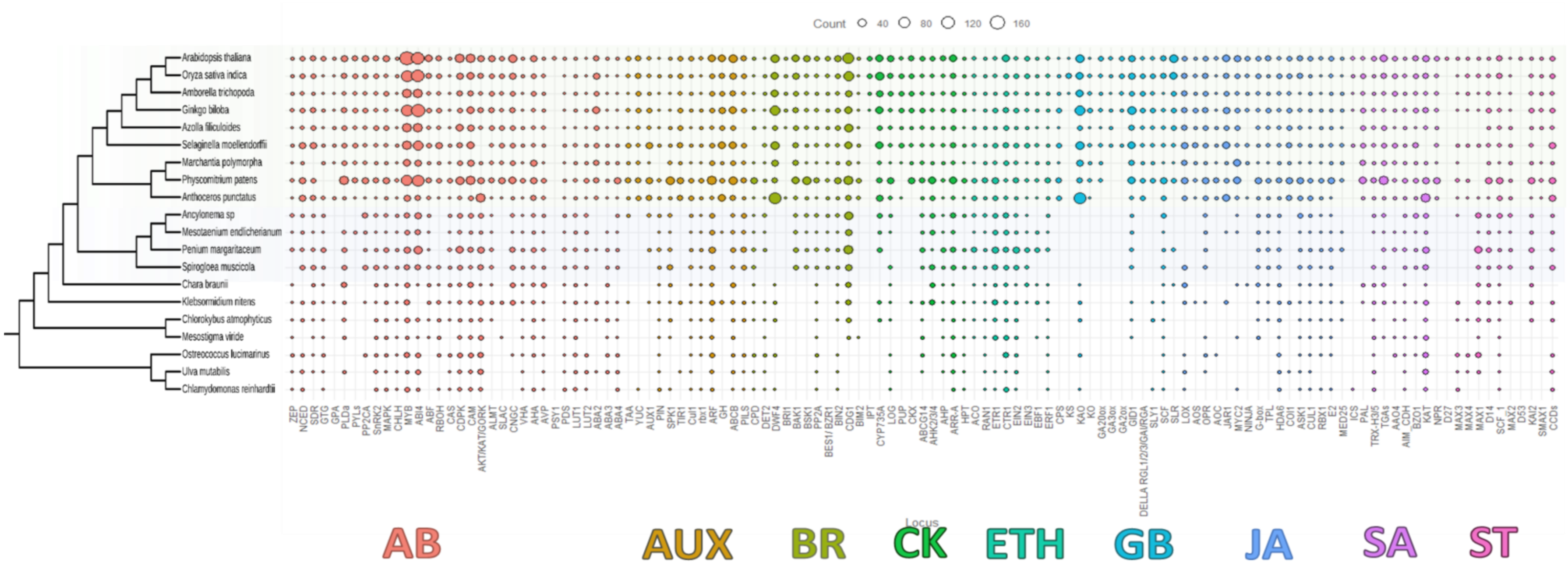


**Supplementary Figure 18** The evolution of phytohormone signalling. Gene count data for each species in the comparative genomic dataset for genes involved in Abscisic Acid (ABA), Auxin, Brassinosteroids (BR), Cytokinins (CK), Ethylene (ETH), Gibberellic Acid (GB), Jasmonic Acid (JA), Salicyclic Acid (SA) and Strigolactones (ST).

### Supplementary Information 8

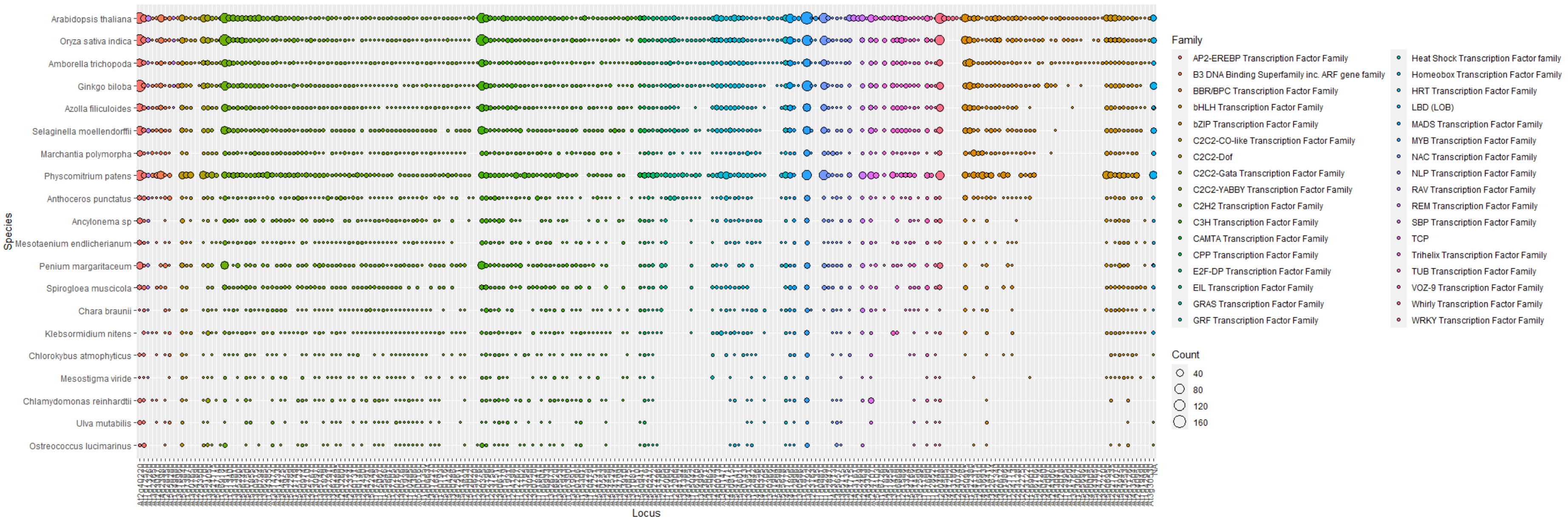


**Supplementary Figure 19** The evolution of transcription factors. Gene count data for each species in the comparative genomic dataset for AP2-EREBP, B3 DNA Binding, BBR/BPC, bHLH, bZIP, C2C2-CO-like, C2C2-Doc, C2C2-Gata, C2C2-YABBY, C2H2, C3H, CAMTA, CPP, E2F-DP, EIL, GRAS, GRF, Heat shock, Homeobox, HRT, LBD, MADS, MYB, NAC, NLP, RAV, REM, SBP, TCP, Trihelix, TUB, VOZ-9, Whirly and WRKY Transcription factor families.

**Supplementary Information 9**

**Fossil calibrations for Molecular Clock Analysis**

Phylogenetically relevant calibrations were taken from those used in Bowles et al (2023)[21] and are briefly discussed below.

**Calibration_1:** Viridiplantae/Chloroplastida: Chlorophyta - Streptophyta

**Fossil taxon and specimen:** *Proterocladus antiquus* [27]
***Phylogenetic justification:*** Following Harris *et al.* [28]
***Minimum age:*** 940.4 Ma.
***Soft maximum age:*** 1879.6 Ma.
***Age justification:*** Following Harris *et al.* [28]

**Calibration_2:** Chlorophyta

**Fossil taxon and specimen:** *Proterocladus antiquus* [27]
***Phylogenetic justification:*** Following Harris *et al.* [28]
***Minimum age:*** 940.4 Ma.
***Soft maximum age:*** 1879.6 Ma.
***Age justification:*** Following Harris *et al.* [28]

**Calibration_3:** Chlorophyceae – Ulvophyceae

***Fossil taxon and specimen***: *Palaeocymopolia silurica* [29]
***Phylogenetic justification:*** Following Morris et al [30]
***Minimum age:*** 438.3 Ma.
***Soft maximum age:*** 1879.6 Ma.
***Age justification:*** Following Morris et al [30]

**Calibration_4:** Trebouxiophyceae

**Fossil taxon and specimen:** *Botryococcus sp (previously Gloeocapsomorpha sp)* from the Upper Permian Hyland Bay Formation, Western Australia [31].
***Phylogenetic justification:*** Extant *Botryoccocus* are planktonic colonial algae found in brackish and fresh water. Their outer cell walls are formed of an acid-resistant biopolymer. Colonies of fossilised *Botryococcus* consist of cell clusters, with individual cells being oblong [31].
***Minimum age:*** 298.75 Ma.
***Soft maximum age:*** 1879.6 Ma Ma.
***Age justification:*** Botryococcus appears early within the fossil record, dating from the Precambrian (>542Ma) [32–35]. They are the largest contribution to the formation of crude oil and oil shales. In oil shales botryococcanes are present, which are lipids that co-localise with fossil Botryococcus [36].

**Calibration_5:** Streptophyta: Charophyta – Embryophyta

***Fossil taxon and specimen***: *Tetrahedraletes* cf. *Medinensis* [37].
***Phylogenetic justification:*** Following Morris *et al.* [30]
***Minimum age:*** 469 Ma.
***Soft maximum age:*** 1879.6 Ma Ma.
***Age justification:*** Following Morris *et al.* [30]

**Calibration_6**: Embryophyta: Bryophyta – Tracheophyta

***Fossil taxon and specimen:*** *Tetrahedraletes cf. medinensis*
***Phylogenetic justification:*** Following Morris *et al.* [30]
***Minimum age:*** 469 Ma
***Maximum age:*** 515.5 Ma
***Age justification:*** Following Morris *et al.* [30]

**Calibration_7**: Tracheophyta: Lycopodiophyta – Euphyllophyta

***Fossil taxon and specimen:*** Zosterophyllum sp. and Chelinospora? sp.
***Phylogenetic justification:*** Following Harris *et al.* [28]
***Minimum age:*** 420.7 Ma
***Maximum age:*** 458.88 Ma
***Age justification:*** Following Harris *et al.* [28]

**Calibration_8**: Euphyllophyta

***Fossil taxon and specimen:*** *Kenrickia bivena*
***Phylogenetic justification:*** Following Harris *et al.* [28]
***Minimum age:*** 393.2 Ma
***Maximum age:*** 451 Ma
***Age justification:*** Following Harris *et al.* [28]

**Calibration_9**: Spermatophyta: Acrogymnospermae – Angiospermae

***Fossil taxon and specimen:*** *Cordaixylon iowensis*
***Phylogenetic justification:*** Following Morris *et al.* [30]
***Minimum age:*** 308.14 Ma
***Maximum age:*** 365.63 Ma
***Age justification:*** Following Morris *et al.* [30]
